# Live attenuated YFV-17D vaccine induces robust B and T cell immunity in vaccine-draining lymph nodes

**DOI:** 10.64898/2026.11.04.764404

**Authors:** Rachel B Evans, Wen Shi Lee, Palak Mehta, Christopher A Gonelli, Ming ZM Zheng, Robyn Esterbauer, Ellie Reilly, Kathleen M Wragg, Andrew Kelly, John J Zaunders, Tri Giang Phan, Summa Bibby, Naphak Modhiran, Daniel Watterson, Solange Obeid, Brad Milner, Stephen J Kent, Anthony D Kelleher, C Mee Ling Munier, Jennifer A Juno, Adam K Wheatley

**Affiliations:** Department of Microbiology and Immunology, University of Melbourne, at The Peter Doherty Institute for Infection and Immunity, Melbourne, Victoria, Australia; Immunovirology and Pathogenesis Program, Kirby Institute, UNSW Sydney, NSW, Australia; St Vincent’s Hospital Sydney, Darlinghurst, NSW, Australia; Precision Immunology Program, Garvan Institute of Medical Research, Sydney, NSW, Australia; School of Chemistry and Molecular Biosciences, The University of Queensland, Brisbane, Queensland, Australia; Australian Infectious Disease Research Centre, Global Virus Network Centre of Excellence, Brisbane, Queensland, Australia; Melbourne Sexual Health Centre and Department of Infectious Diseases, Alfred Hospital and Central Clinical School, Monash University, Melbourne, VIC, Australia

**Keywords:** FNA, FNB, YFV-17D, lymph node, yellow fever virus, TFH, germinal centre

## Abstract

The yellow fever virus-17D (YFV-17D) vaccine is widely regarded as one of the most effective vaccines in existence, with protection linked to robust elicitation of neutralising antibodies^1^. The features of the germinal centre (GC) reaction driving this durable and potent antibody responses remain unclear. Here we examined immune dynamics in blood and lymph nodes (LNs) after primary YFV-17D immunisation in humans. Using fine needle biopsy (FNB), we identified robust T and B lymphocyte responses within draining LNs, underpinning maturation of antibodies targeting the viral envelope (E). Early antibody responses were largely IgM, with gradual accumulation of E-specific memory B cells and anti-viral IgG in the subsequent months. Monoclonal antibodies derived from GC B cells had specificity for E, with diversification in the LN capable of driving increases in binding and neutralising potency. While circulating vaccine-specific CD4^+^ T cells peaked two weeks post-vaccination, clonally related GC and IL-10^+^ T follicular helper populations progressively accumulated in the LN. Comparison of FNB transcriptional datasets following YFV-17D, SARS-CoV-2 mRNA/LNP, and seasonal influenza vaccination identified a conserved GC program involving similar B and T lymphocyte subpopulations. Collectively, we find that while YFV-17D induces robust GC activity, this trait is shared with other vaccine platforms and the notably durable protection against infection may reflect an incapacity of the virus to escape host immunity.

## Main

Vaccines are a critical tool for reducing the burden of infectious diseases. The live-attenuated yellow fever virus strain 17D (YFV-17D) vaccine, with hundreds of millions of doses administered and a proven ability to elicit durable protective immunity, stands out as a “gold-standard” to inform the development of effective vaccines still lacking for many critical pathogens. Immunity is typically generated within secondary lymphoid tissues draining the vaccine administration site. These lymphoid tissues house germinal centre (GC) structures that facilitate critical interactions between T and B lymphocytes, which are essential for generation of high-quality antibodies. We therefore initiated a longitudinal study of 17 healthy adults who received primary YFV-17D (Stamaril) immunisation via intramuscular injection, with regular collection of blood, and fine needle biopsy (FNB) of the draining axillary lymph nodes (LNs) from 6 participants taken at weeks 2 (n=3) and 4 (n=3) post-immunisation (**Fig. 1a**; **Supplementary Table 1**).

**Figure 1.**
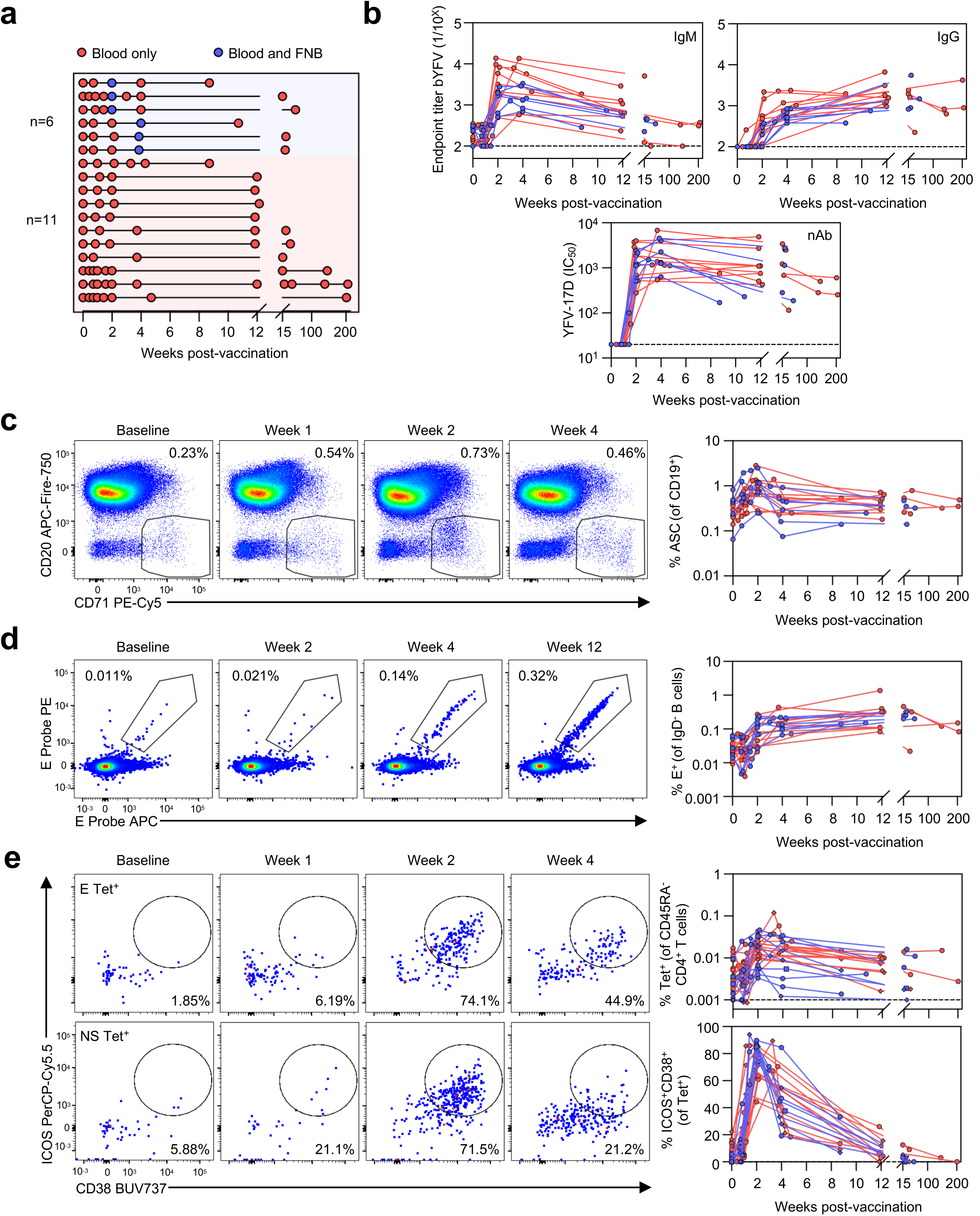
Immune Dynamics of YFV-17D immunization in the blood. **(a)** Study design and longitudinal sampling of each participant. **(b)** Serological outcomes of primary YFV-17D vaccination measured by Binjari YFV (bYFV) virus-like particle IgM (top left) or IgG (top right) ELISA or live virus neutralization (bottom). **(c)** Representative staining and frequency of ASCs (CD20^lo^ CD71^+^). **(d)** Representative staining and frequency of E probe^+^ cells among IgD^-^ B cells (CD19^+^CD20^+^). **(e)** Representative staining of ICOS and CD38 on pHLA-DR tetramer^+^ (Tet^+^) cells specific for E (top left) and NS epitopes (bottom left). Graphs indicate frequency of Tet^+^ cells (top right), and proportion of ICOS^+^CD38^+^ Tet^+^ cells (bottom right). Symbols indicate tetramer specificity for E (circles), C (squares) or NS epitopes (diamonds). (a-d), N=17; (e) N=14. Red, participants with blood sampling only; blue, participants with blood and FNB sampling.

### Dynamics of the peripheral immune response to YFV-17D

To establish the kinetics of the response to YFV-17D in blood, we undertook serological analysis of antibodies within plasma, and flow cytometric analysis of PBMCs. Early post-vaccination, all subjects displayed a strong induction of neutralising antibodies (nAbs), YFV-specific IgM, and antibody secreting cells (ASCs) (**Fig. 1b, 1c**). By contrast, YFV-specific IgG exhibited delayed kinetics, slowly increasing over the months following vaccination (**Fig. 1b**). nAbs strongly correlated with YFV-specific IgM titres in the early weeks post-vaccination but correlated with IgG titres at later timepoints (**Extended Data Fig. 1a**). Viral envelope (E)-specific memory B cells (B_MEM_) rapidly became activated **(Extended Data Fig. 1b)** and increased in frequency, peaking in the blood approximately 12 weeks post-vaccination (**Fig. 1d**). The isotype dynamics of E^+^ B_MEM_ cells aligned with serological data (**Extended Data Fig. 1c**), with IgM^+^ cells peaking early. Frequencies and activation of circulating CD4^+^ T cells specific for E, capsid (C) and non-structural (NS) peptides generally peaked around week 2 post-vaccination (**Fig. 1e, Extended Data Fig. 1d**) and predominantly exhibited CXCR5^-^PD-1^+^ and CCR6^-^CXCR3^+^ phenotypes (**Extended Data Fig. 1e,f**).

Following immunisation, YFV-17D RNA can be readily detected in the blood of vaccine recipients^2,3^, raising the question of whether the immune response is initiated in the LNs or is more broadly disseminated. We first turned to a murine model whereby C57BL/6 mice were vaccinated intramuscularly with 10^4^ PFU of YFV-17D (**Extended Data Fig. 2a**). Robust induction of nAb was observed in the serum by 2 weeks post-immunisation (**Extended Data Fig. 2b**), in line with the human cohort. Total and E-specific GC B cell responses were concentrated to the LNs draining the vaccination site, rather than the spleen and distally located non-draining LNs (**Extended Data Fig. 2c,d**). Likewise, T_FH_ cells were predominantly found in the draining LNs at week 2 post-vaccination (**Extended Data Fig. 2e,f**). Given the LN-localised nature of the murine response, we sampled the draining axillary LNs after YFV-17D vaccination in our clinical cohort via FNB.

### Biogenesis of YFV-17D lymph node germinal centre reaction

To enable in-depth transcriptional characterisation of vaccine-specific B and CD4^+^ T cells in FNB of lateral axillary LN, we used flow cytometry to sort populations of B and T cells that were not naïve (IgD^-^ or CD45RA^-^) and were either expressing canonical activation markers and/or were antigen-specific (based on binding to E probes or a cocktail of HLA-matched E, C, NS1 and NS3 pHLA tetramers, see Methods for details of alleles and epitopes) (**Extended Data Fig. 3a,b**). Contralateral LN samples collected from 2 participants confirmed enrichment of antigen-specific GCs in the vaccine draining LN (**Extended Data Fig. 3c**). Single cell gene expression, T cell receptor (TCR)/B cell receptor (BCR) sequencing, and antibody derived tag (ADT) analysis was performed for a total of 22149 IgD^-^ B cells and 29706 CD45RA^-^ CD4^+^ T cells, with each participant and tissue distinguished using hashtags. The resulting dataset resolved 11 transcriptionally distinct B cell clusters and 17 CD4^+^ T cell clusters distributed across blood and LN (**Extended Data Fig. 4a,c; Supplementary Table 2**) that were largely independent of cell cycle (**Extended Data Fig. 4b,d**).

Unbiased clustering of single B cell transcriptional signatures identified canonical immune cell populations (**Fig. 2a**) with gene expression profiles consistent with prior reports^4,5^. Prominent populations of GC B cells (clusters f, g, h) largely restricted to LNs **(Extended Data Fig. 4a**), expressed *BCL6, GCSAM* and *S1PR2* genes and surface markers CD10, CD95 and CD38 (**Fig. 2b**). Antibody-secreting or plasma cells (cluster i) were observed in both LN and PBMC and expressed high levels of canonical markers *PRDM1* and *IRF4*. Cluster d delineated an activated, class-switched B cell population, and additionally a small population of B cells responsive to interferon signalling (cluster e) was observed. Three populations of resting B_MEM_ cells were distinguished by IgM-(cluster a), IgG-(cluster b) and T_H_2-bias (cluster c)^6,7^. GCs were already observed within the draining LN at week 2 post immunisation (**Extended Data Fig. 5a**). Limited B cell binding to E probes was observed at this timepoint, although there was evidence of clonal expansion (**Extended Data 5b,c**) and expression of genes such as *S1PR2* and *BCL6* consistent with GC activity (**Extended Data Fig. 5d**). By 4 weeks post-immunisation, robust clonally expanded populations of E and non-E specific GC B cells could be observed in the three vaccine recipients sampled at this timepoint (**Fig. 2c**), with some families exceeding 200 individual clones (**Fig. 2d; Supplementary Table 3**). The frequency of somatic mutation within immunoglobulin heavy chain variable regions (*IGHV*) was generally low (<5%), and most clonal families predominantly utilised either *IGHM* or *IGHG1* constant chains.

**Figure 2.**
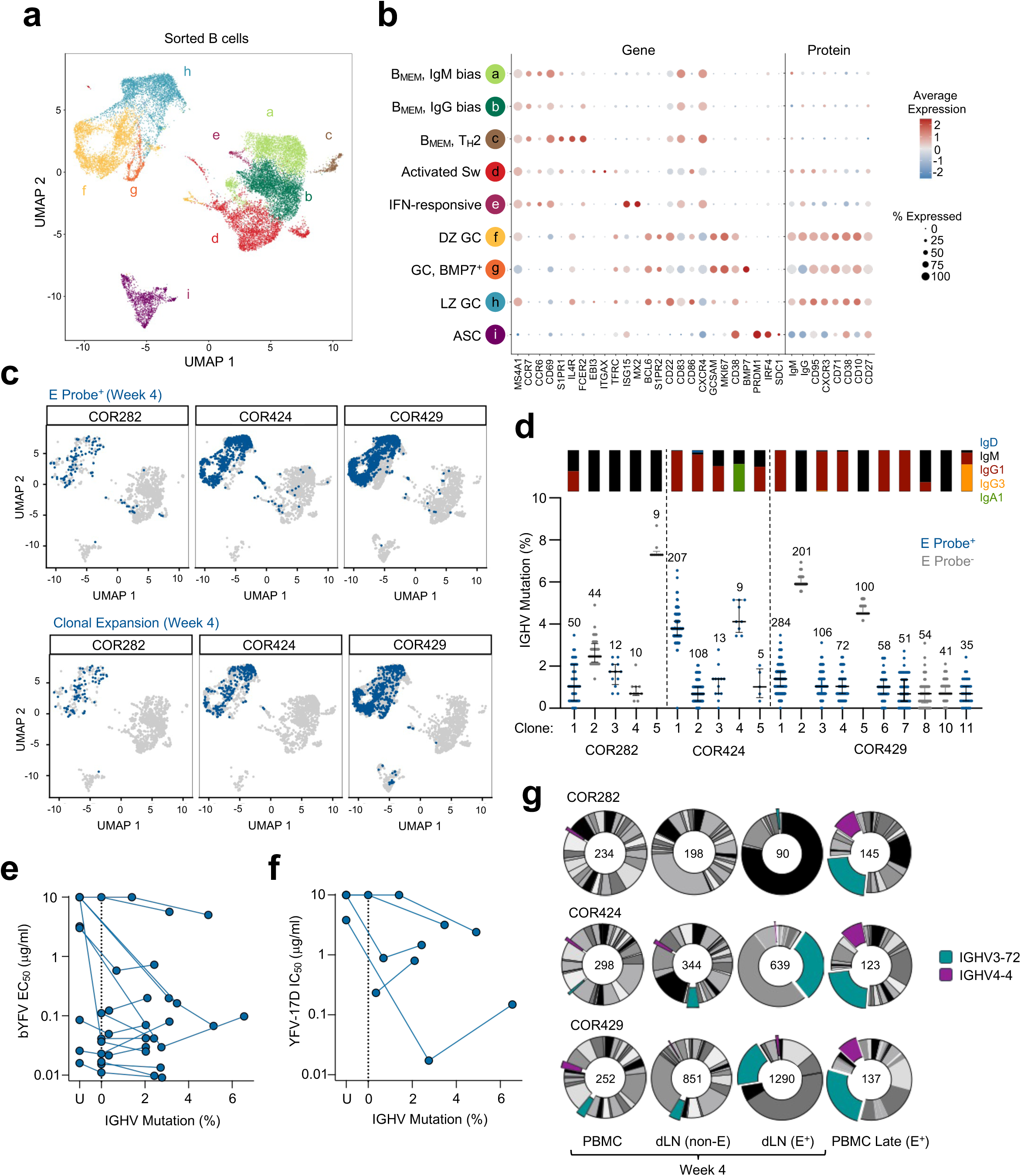
Transcriptional and immunoglobulin profile of YFV-specific B cells in the blood and draining lymph node. **(a)** UMAP clustering of all sorted B cells from blood and dLN at weeks 2 (N=3) and 4 (N=3). **(b)** Expression of key genes and proteins among selected B cell clusters. **(c)** UMAP overlay of E Probe^+^ or clonally expanded B cells (blue) identified at week 4 post-immunization for participants COR282, COR424 and COR429. **(d)** Immunoglobulin sub-class and *IGHV* nucleotide mutation frequency of selected clonally expanded B cell families (median and IQR). **(e)** ELISA binding of recombinant mAbs or inferred unmutated common ancestors (U) to binjari YFV (bYFV) virus-like particles. (**f**) Neutralisation activity of mAbs against live YFV-17D virus. **(g)** IGHV gene utilisation from sorted PBMC or LN B cell populations at week 4 post immunisation, or from E-specific memory B cells subsequently recovered within PBMC at a late timepoint (weeks 13-25). Proportions of IGHV3-72 and IGHV4-4 use are highlighted.

To examine the specificity of GC B cells, we generated recombinant monoclonal antibodies (mAbs) from representative clonal families for each participant. Thirty mAbs were generated from GC B cells at week 2 post-YFV-17D, however none showed binding to YFV-17D E or viral neutralisation (**Supplementary Table 4**). In contrast, multiple antibody lineages isolated by 4 weeks post-immunisation demonstrated binding (**Fig. 2e**) and/or neutralisation activity (**Fig. 2f**), which generally matured in line with the acquisition of somatic mutations.

To better understand the fate of E-specific B cells, we sequenced E-specific B_MEM_ cells in the blood from later timepoints (weeks 9-53 post-immunisation). Consistent with a previous study^8^, we found enrichment within E-specific immunoglobulins of a stereotypic class of IGHV3-72-utilising antibodies reportedly directed to the fusion loop domain. Additionally, we see potential enrichment of antibodies utilising IGHV4-4 (generally paired with IGLV1-51), which are over-represented within nAb and are known to target epitopes within domain II of E overlapping the prototypical mAb 5A^9^ (**Fig. 2g, Extended Data 5e**). Within FNB samples, we found IGHV3-72 enrichment evident at 4 weeks but not 2 weeks. Overall, these data suggest that GC activity observed 2 weeks post-immunisation may reflect residual activity from prior antigenic encounters rather than the nascent response to YFV-17D.

Similar to the B cell response, profiling of CD4^+^ T cells identified canonical GC and T_H_1/T_H_2/T_H_17 populations. Clusters were broadly defined as ‘resting’ (high *IL7R*, low ICOS and CD38; clusters a-e) or ‘activated’ (low *IL7R*, high ICOS and CD38; clusters f-n) (**Fig 3a, Extended Data Fig. 4c**). Cluster e was enriched in the PBMC dataset and expressed moderate levels of *CXCR5*, consistent with a circulating T_FH_ cell phenotype. Cluster j was distinguished by expression of interferon-related transcripts, while cluster o (‘T_REG_’) was clearly defined by *FOXP3, IL2RA* and *IKZF2* expression. Canonical GC T_FH_ cells (cluster m) expressed the highest levels of *BCL6, CXCR5* and *PDCD1.* Clusters with moderate *BCL6* expression included clusters l (high *MKI67*, ‘proliferating’), k (‘pre-T_FH_’, intermediate *CXCR5* and *PDCD1*), and n (*IL21* and *IL10* expression, ‘IL-10^+^ T_FH_’^10,11^) (**Fig 3b, Supplementary Table 2**).

**Figure 3.**
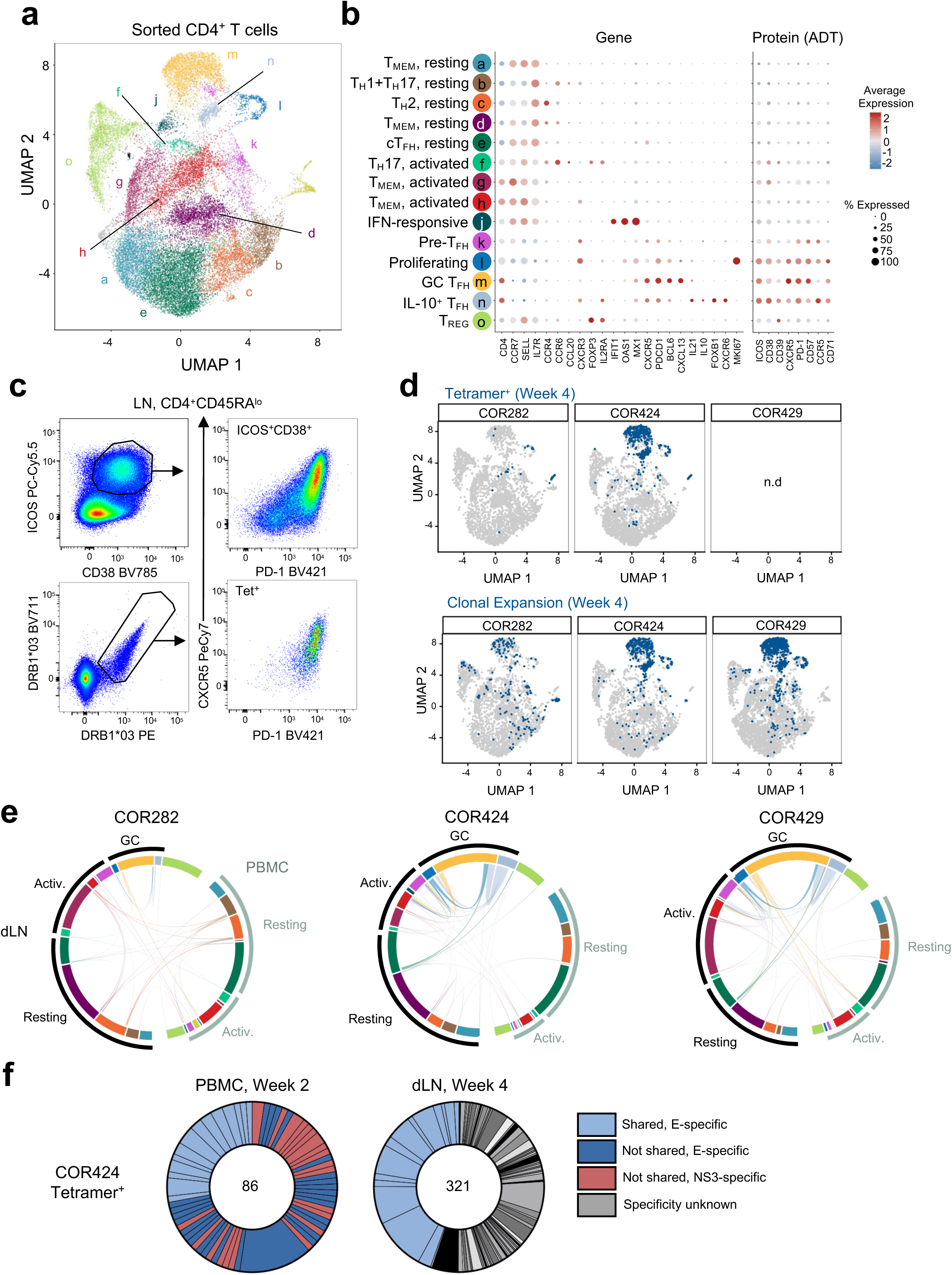
Transcriptional profile and clonal overlap of YFV-specific CD4^+^ T cells in the blood and draining lymph node. **(a)** UMAP clustering of all sorted CD4^+^ T cells from blood and dLN at weeks 2 (N=3) and 4 (N=3). **(b)** Expression of key genes and proteins among selected CD4^+^ T cell clusters. **(c)** Frequency and phenotype of tetramer^+^ CD4^+^ T cells in PBMC and dLN for COR424. **(d)** Overlay of tetramer^+^ or clonally expanded T cells (blue) at week 4 for participants COR424 and COR429. **(e)** Circos plots for each week 4 participant showing shared and unique clonotypes between transcriptional clusters in the dLN and PBMC. **(f)** Distribution of TCR clonotype antigen specificity among Tet^+^ cells recovered from sorted PBMC (week 2) or dLN T cells (week 4) for COR424.

All CD4^+^ T cell clusters were present in the draining LN at week 2, including the IFN-responsive cluster that was largely absent at week 4 (**Extended Data Fig. 6a**). Consistent with the B cell dynamics, both GC and IL-10^+^ T_FH_ cell populations were evident at week 2 post-vaccination and were higher in frequency by week 4 (**Extended Data Fig. 6a**). pHLA tetramer^+^ cells in the draining LN were rare at week 2 (**Extended Data Fig. 6b,c**) but robustly detected at week 4, where they predominantly exhibited a T_FH_ cell phenotype by both flow cytometry (**Fig 3c**) and transcriptional profiling (**Fig 3d**). At both timepoints, the T_FH_ cell clusters (l, m, n) contained most clonally expanded T cells, with extensive clonotype sharing across clusters (**Fig 3e, Extended Data Fig. 6e; Supplementary Table 5**). Matching clonotypes were identified across blood and draining LN samples for all participants, particularly between GC T_FH_ cells and an activated memory T cell PBMC cluster (cluster h).

Given the robust population of E and NS3 tetramer^+^ T_FH_ cells at week 4 for COR424, we sorted and performed TCR sequencing of E-specific or NS3-specific tetramer^+^ cells from PBMC at 2 weeks post-vaccination. 27.9% (24/86) of the TCRs within PBMC were clonally related to tetramer^+^ cells in the draining LN, and all were E-specific (**Fig. 3f**). Collectively, these clonotypes accounted for 44.5% of the tetramer^+^ cells found in the draining LN at week 4, indicating a shared clonal origin of effector/memory T cells exported into the circulation and T_FH_ cells found in the GC two weeks later.

### Conservation of GC transcriptional profiles across multiple vaccines

To assess whether YFV-17D induces a transcriptionally unique GC response, we compiled available data from FNB studies of seasonal influenza^10,12,13^, adjuvanted influenza^14^, SARS-CoV-2 mRNA^11,15,16^, and influenza mRNA vaccines^17^ at nascent/pre-GC (∼2 weeks) and established GC (∼4 weeks) timepoints. Naïve lymphocyte populations were excluded for consistency with the present study. The B cell and T cell transcriptional clusters from the YFV-17D samples (**Extended Data Fig 4a, c**) were mapped onto each dataset to identify cells with a similar GEX profile. Despite substantial differences in vaccine platform (live-attenuated, mRNA/lipid nanoparticle, inactivated virus) and participant immune history (primary single-dose, prime/boost, and pre-existing memory), B cell and CD4^+^ T cell subpopulations similar to those present following YFV-17D were identified across FNB datasets (**Fig. 4a-b, Extended Data Fig. 7a-b**). While direct quantification of cluster prevalence is challenging due to methodological differences between studies, expression patterns of key genes suggest that core transcriptional GC programs are maintained regardless of vaccine product (**Fig 4c-d, Extended Data Fig 7c-d**).

**Figure 4.**
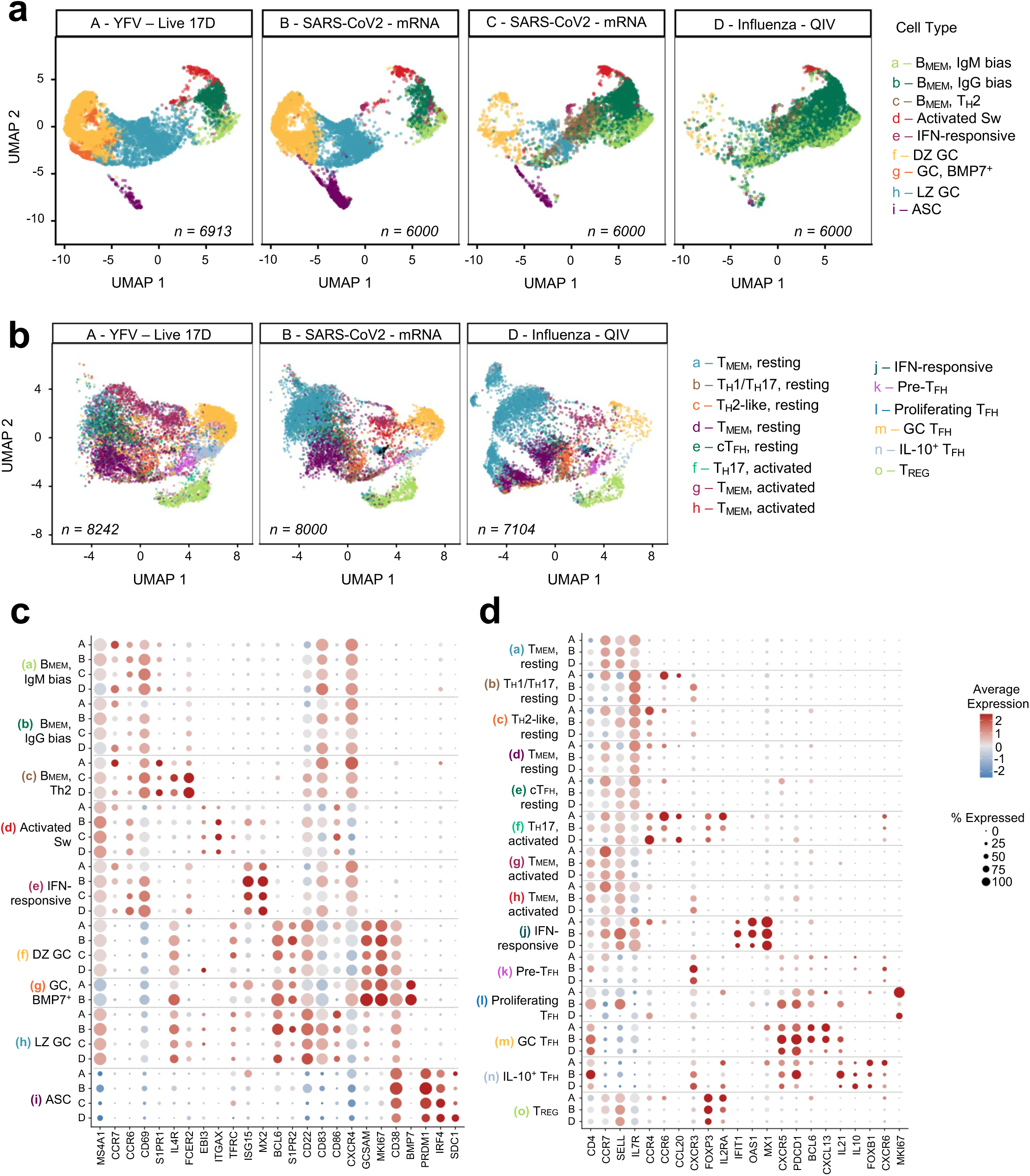
Transcriptional profile of lymph node B and T lymphocyte populations across multiple vaccine platforms. UMAP representations of lymph node **(a)** B cell and **(b)** CD4^+^ T cell populations observed after immunization with YFV-17D (A; this study), SARS-CoV-2 mRNA vaccines (B; Kim et al.^15^ for B cells, Schattgen et al. for T cells^10^, C; Dhenni et al.^16^) or seasonal influenza vaccines (D; McIntire et al.^12^). Following dataset integration of the different studies, predicted YFV-17D cluster identities from Figure 2A or 3A were projected onto a shared UMAP. Dotplots detailing expression of key genes among **(c)** B cell or **(d)** CD4^+^ T cell clusters for each dataset.

In addition to FNBs, tonsils serve as a relatively accessible secondary lymphoid tissue for the study of human GCs. B cell and CD4^+^ T cell gene expression data was extracted from a human tonsil atlas^18^ where adults underwent tonsillectomy for sleep apnea or tonsillitis. Overall, the GC B cell and T_FH_ cell populations identified in the YFV-17D LN were present in the tonsil datasets (**Extended Data Fig. 8a,b**) and expressed canonical, lineage-defining genes including *BCL6, GCSAM, PRDM1*, and *FOXP3* (**Extended Data Fig. 8c,d**). Nevertheless, several tissue-specific differences were observed, including greater *IGHA1* and *FCRL5* in tonsillar B cells but lower expression of genes involved in T-B interactions such as *BACH2, CXCL13* and *SH2D1A* (SAP) (**Extended Data Fig 8e, f**), suggesting potential modulation of GC activity in chronically inflamed tissues.

## Discussion

The live-attenuated YFV-17D is notable for its high efficacy, with protective nAbs induced in the vast majority (<95%) of recipients after a single dose^19^. Within our cohort, YFV-17D induced robust serum nAbs, which arose 2 weeks post-vaccination and preceded the establishment of E-specific GC activity in the draining LNs. The potency of the early nAb response appears to be a relatively unique feature of YFV-17D and contrasts with the multiple vaccine doses often required to elicit maximal nAb activity against viruses such as SARS-CoV-2^20^, hepatitis B^21^, or Nipah^22^. Here, early nAb titres were strongly correlated with YFV-specific IgM in agreement with past findings^8,23^. Initial nAb emergence temporally aligned with peak ASC frequencies, consistent with the previously reported high proportion of early ASC-derived mAbs exhibiting potent neutralization^8^.

The notable potency of the early IgM nAb response suggests that the unusually high efficacy of YFV-17D might partly reflect an intrinsic susceptibility of orthoflaviviruses to neutralisation^24,25^. Combined with a comparative lack of extensive viral sequence or antigenic evolution^26–28^, these factors could markedly impact the apparent durability of vaccine-induced protection. Similarly, the existence of multiple public stereotypic pathways for antibody targeting of the viral E protein^8,29^ may provide a unique jump start in the genesis of host humoral immune responses against YFV-17D, which may not be evident for other pathogens.

By week 4, expansion of E^+^ class-switched B cells with a classical GC phenotype was observed in the draining LN, while circulating E^+^ B_MEM_ cells slowly accumulated in the blood over several months post-vaccination. Antibodies derived from E^+^ GC B cells showed evidence of affinity maturation and gains in neutralization potency, and we confirmed enrichment of IGHV3-72-derived immunoglobulins corresponding to a stereotypic class known to target the fusion loop^8^. The continued emergence of E-specific, class-switched and somatically mutated B_MEM_ cells into blood is suggestive of ongoing GC activity for several months after vaccination, in line with the durability of GCs reported for other vaccines in humans^4,30^ and nonhuman primates^31,32^. The rapid clearance of circulating virus and reliable induction of neutralising IgM suggests GC-activity induced by YFV-17D primarily serves as an engine for diversification of immunological memory and is not strictly required for direct antiviral function.

In contrast with the protracted nature of the E-specific B cell response, YFV-17D tetramer-specific CD4^+^ T cells showed peak activation and expansion in the blood by 2 weeks post-vaccination, consistent with other studies^33–35^. Indeed, tetramer^+^ cells largely returned to a resting phenotype by the time vaccine-specific GC activity in the LN was established at week 4. Nevertheless, we observed extensive clonal overlap between LN T_FH_ cell populations and circulating CD4^+^ T cells in both time-matched and longitudinal samples, despite the low frequency of CXCR5^+^ tetramer^+^ cells detected in the blood. Similarly discordant kinetics and phenotypes between the blood and lymphoid compartment have also been observed following SARS-CoV-2 mRNA vaccination^30^, highlighting the challenge of identifying circulating CD4^+^ T cell populations that correlate with ongoing GC activity. Within the LN, YFV-specific CD4^+^ T cells differentiated into both prototypical GC T_FH_ cells and a clonally linked IL-10^+^ T_FH_ cell population that has been consistently reported in other immune contexts^10,11,32^ but whose function is currently not well defined.

The significant logistical challenges in directly sampling human LN tissues have hampered our understanding of human vaccine immunology, with clinical studies typically limited to peripheral blood. While methodological differences between studies complicates direct comparison of cell proportions, we observe substantial commonalties in the composition and gene expression profiles of canonical LN B and T lymphocyte sub-populations between YFV-17D and other vaccine platforms, particularly primary mRNA immunisation for SARS-CoV2^15,16^. Minor populations with poorly defined functions in the LN, including BMP7+ GC B cells and IFN-responsive cells, may represent distinct features of specific vaccine platforms or transient temporal changes in GC state. Nonetheless, shared transcriptional programming in LN populations, which extend to human tonsil samples, suggest that GCs may follow a relatively conserved trajectory once formed, in line with cross-species conservation of GC structure and evolutionary constraints on optimal environments for T-B cell interaction^36^. Future studies allowing direct experimental comparison of the immunogenicity of different vaccine technologies in longitudinal LN and blood samples will enable finer dissection of optimal GC characteristics for maximising protective immunity.

## Methods

### Participant recruitment, human ethics and sample collection

Study protocols were approved by human research ethics committees for the University of Melbourne (20314 and 27049) and or St Vincent’s Hospital Sydney (2023/ETH00180). All participants provided written informed consent in accordance with the Declaration of Helsinki.

17 healthy participants were enrolled and received intramuscular immunisation with the live-attenuated YFV-17D (Stamaril) vaccine. Peripheral blood samples were collected in heparin and serum tubes. Plasma and serum were collected and frozen at -80C. Peripheral blood mononuclear cells (PBMC) and granulocytes were isolated by Ficoll-Paque separation. PBMC were washed, cryopreserved in 10% DMSO/90% fetal calf serum (FCS), and stored in liquid nitrogen. DNA was extracted from granulocytes using the QIAamp DNA Micro Kit (Qiagen) for HLA typing (Australian Red Cross). Six of the vaccine cohort participants (3 male, 3 female, aged 24-43 years old) additionally underwent axillary LN FNBs at days 14-28 post-vaccination. FNBs were performed by a qualified radiologist and ultrasonographer at St Vincent’s Hospital, Sydney as previously described^37^. Briefly, axillary LNs were identified by ultrasound, local anaesthetic was administered, and cells were collected by four passes of a 25-gauge needle in each LN. Following each pass, cells were transferred into a 15 mL tube. RF10 media was drawn into the syringe and expelled to wash the needle of any residual cells. Cells from an ipsilateral (vaccine-draining) LN were successfully collected from 5 of 6 participants, with two contralateral LN samples also obtained. Peripheral blood was collected from all individuals. LN cell suspensions from COR282, 424, 429 and 430 were additionally subject to red blood cell lysis for 3 minutes at room temperature (BC Pharm Lyse, 1:10 diluted in dH_2_O) prior to immunophenotyping and cell sorting. Paired PBMC samples were isolated by Ficoll gradient centrifugation as described above. 1×10^7^ PBMC were freshly used for immunophenotyping and sorting, with the remainder cryopreserved.

### Animal Studies

C57BL/6 mice were bred in-house under specific pathogen-free conditions in the animal facility at the Peter Doherty Institute of Infection and Immunity, University of Melbourne, Australia. Mouse studies were carried out in accordance with the University of Melbourne Animal Ethics Committee (approval no. 30633 and 32601). All mice were female and aged 6–12 weeks at the time experiments commenced. For YFV-17D vaccination, 10^4^ PFU of virus was prepared and mice were anaesthetised by isoflurane inhalation with oxygen flow at 2 L/min and isoflurane vapouriser set to 3 (Stinger Anaesthetic Machine). Virus was delivered in 50 μl intramuscular injections at the left and right quadriceps using a 29G needle. At experimental endpoints, mice were killed using CO_2_ asphyxiation using a delivery of 50% chamber volume per minute.

### Expression of YFV-17D antigens

Nucleotide sequences encoding the YFV-17D E protein (ENA X03700.1, AA286-677) were synthesised with an N-terminal prolactin signal sequence and C-terminal AviTag and polyhistidine tag and cloned into pFastBac before being used to generate baculovirus vector using the Bac-to-Bac baculovirus system with ExpiSf cells according to the manufacturer (ThermoFisher Scientific). Recombinant E protein was produced in ExpiSf cells treated with ExpiSf Enhancer (ThermoFisher Scientific) 18– 24hr pre-infection and using 5–10 MOI baculovirus for infection. Clarified culture supernatants were collected 3 days post infection and E protein was purified using Ni Sepharose excel resin (Cytiva). The purified E was buffer exchanged into 10mM Tris-HCl, pH 8.0 and biotinylated using BirA ligase (Avidity) before being further purified by size-exclusion chromatography with a Superdex 200pg 16/600 column (Cytiva) to select for dimeric protein. For use as flow cytometric B cell probes, biotinylated E was complexed via serial addition of streptavidin-PE (ThermoFisher Scientific) or streptavidin-APC (ThermoFisher Scientific).

### Production of Binjari YFV-17D virus like particles

Chimeric virus like particles consisting of the prME genes of YFV-17D (GenBank: KF769015.1) and the genetic backbone of Binjari virus (BinJV), were prepared as generated and prepared as previously described^38^. Briely, C6/36 (*Aedes albopictus*) cells were infected with BinJV/YFV-17D (bYFV) at an MOI of 0.01 in RPMI media supplemented with 2% FCS (Bovogen) and 100 U/mL penicillin and 100 μg/mL streptomycin (Gibco). On days 5, 7 and 10 post infection, the supernatant was collected, clarified via centrifugation (15 mins at 3000 x g) and sterilised by passing through a 0.22 μm filter. Virus was precipitated in the pooled supernatant by adding polyethylene glycol 8000 (final concentration 4% w/v) in NTE buffer (120 nM NaCl, 10 mM Tris-Base and 1 mM EDTA, pH 8) and incubating overnight at 4°C before centrifugation at 12,000 x g for 1 hour at 4 °C using an Avanti J-26 XPI JLA-10.500 rotor. The virus-containing pellet was resuspended in NTE buffer and further purified via a 20% w/v sucrose cushion, centrifuged at 133,907 × g for 2 hours at 4 °C using a Beckman Coulter Optima L-100 XP SW32Ti rotor. The virus pellet was resuspended in 500 μL of NTE buffer overnight at 4 °C and further purified via a 25-40% potassium tartrate gradient, centrifuged at 336,840 × g for 1 hour at 4 °C using a Beckman Coulter Optima L-100 XP SW60Ti rotor. Virus bands were harvested, buffer exchanged into NTE and stored at 4 °C until use.

Purified bYFV particles were then inactivated using binary ethylenimine (BEI). Briefly, 0.1M 2-bromoethylamine hydrobromide (BEA) in 0.175M NaOH was incubated at 37 °C for 30 minutes to synthesise BEI. The freshly prepared BEI was then combined with purified bYFV particles at 1 mg/mL in a 1:33 v/v ratio and incubated for 24 hours at 37 °C. 1M sodium thiosulphate at a final 1:33:3.4 (BEI:virus:sodium thiosulphate) v/v ratio was added to the virus and BEI mixture to neutralise the remaining BEI. The inactivated bYFV was stored at 4 °C until use.

### Preparation of antigen-specific pHLA tetramers

The following biotinylated peptide/HLA class II monomers (previously described by ^33–35,39^) were produced by ProImmune: HLA-DRB1*03:01/Env_43-59_, HLA-DRB1*03:01/ Env_331-347_, HLA-DRB1*03:01/NS1_85-101_, HLA-DRB1*03:01/NS3_353-367_, HLA-DRB1*15:01/Env_457-473_, HLA-DRB1*15:01/NS3_145-161_, HLA-DRB1*15:01/NS5_367-383_, HLA-DRB1*11:01/Env_373-389_, HLA-DRB1*07:01/Env_415-431_, HLA-DRB1*04:01/Env_301-317_, and HLA-DRB1*01:01/C_49-65_. Tetramers were produced by incubation with SaV-PE (Invitrogen), SaV-APC (Invitrogen), SaV-BB515 (Waters) or SaV-BV711 (Waters) at a 3:1 molar ratio of pHLA:SaV.

#### ELISA

Antibody reactivity to YFV-17D was determined using ELISA. 96-well Maxisorp plates (Thermo Fisher) were coated overnight at 4°C with 1 μg/mL bYFV. After blocking with 5% skim milk powder and 0.1% Tween20 in PBS, wells of serially diluted mAbs, or mouse or human plasma, were added and incubated for two hours at room temperature. Plates were washed prior to incubation for 1 hour at room temperature with HRP-conjugated secondary antibodies: anti-human IgG (Southern Biotech, 1:40000), anti-human IgM (Jackson ImmunoResearch Laboratories, 1:10000), anti-mouse IgG (SeraCare 1:15000). Plates were washed and developed using TMB substrate (Thermo-Fisher) and read at 450nm. Endpoint titres were calculated as the reciprocal serum dilution giving signal 2x background using a fitted curve (4 parameter log regression).

### YFV-17D neutralisation assay with ELISA-based readout

YFV-17D virus was grown in Vero cells from Stamaril^®^ vaccine vials (Sanofi Pasteur) in DMEM/2% FCS. After 7 days, cell culture supernatants containing virions were harvested, clarified via centrifugation at 500g for 10 min, and stored at -80°C. Titers of YFV-17D viral stocks were determined via an infected cell ELISA. Virus was serially diluted in DMEM/2% FCS and incubated on plated Vero cells for 3 days at 37°C. Supernatants were removed and cells fixed with 4% formaldehyde for 30 minutes at room temperature, before washing with PBS and permeabilization with PBS/0.1% Triton X-100 for 15 minutes. Plates were blocked with blocking buffer (PBS/4% bovine serum albumin/0.1% Tween-20) for 1 hour at room temperature before three washes in PBST (PBS/0.05% Tween-20). Recombinant 4G2 antibody (clone D1-4G2-4-15, WEHI Antibody Facility) at 1µg/ml in dilution buffer (PBS/0.2% Tween-20/0.1% BSA/0.5% NP-40) was added for 1 hour. After six washes in PBST, goat anti-mouse IgG HRP secondary antibody (Seracare #5450-0011) was added at a 1:15000 dilution for 1 hour. Plates were developed with TMB, stopped with sulfuric acid and read at 450nm. Stock concentrations yielding absorbance corresponding to 50% infectivity (ID_50_) were determined using sigmoidal dose response curve fitting using Graphpad Prism.

To measure YFV-17D neutralisation, serial dilutions of heat-inactivated plasma or mAbs were incubated with virus stocks at 50×ID_50_ at 37°C for 1 hour, before addition to Vero cells and culture for 3 days at 37°C. ‘Cells only’ and ‘Virus+Cells’ controls demark 0% and 100% infectivity respectively. Plates were developed as above. Absorbance was used to calculate %neutralisation using the formula: (‘Virus+Cells’ – ‘Sample’) ÷ (‘Virus+Cells’ – ‘Cells only’) × 100. IC_50_ values were determined using four-parameter non-linear regression in GraphPad Prism with curve fits constrained between 0% and 100%.

### Immunophenotyping of B and T cells – Human

Cryopreserved PBMC from the YFV-17D vaccine cohort were thawed and stained with antigen-specific B cell probes and surface antibodies. Analysis of B cells was performed with the following antibodies (details in **Supplementary Table 6**): CD11c RB613, IgA VioBlue, CD14 BV510, CD3 BV510, CD8a BV510, CD16 BV510, CD27 BV605, IgG BV786, CD38 AlexaFluor700, CD20 APC-Fire750, CD19 ECD, CD71 PeCy5, IgD PeCy7, IgM BUV395, CD21 BUV737, and Live/Dead Aqua. PBMC were stained with Live/Dead amine reactive dye for 3 minutes at room temperature, quenched with FACS buffer (PBS + 2% fetal calf serum), incubated with E protein probes and surface antibody cocktail for 30 minutes at 4C, washed twice in FACS buffer (spinning at 500xg for 7 minutes), and fixed in 1% paraformaldehyde prior to acquisition on a BD Symphony SE flow cytometer.

Analysis of T cells for longitudinal PBMC analysis was performed with the following antibodies (details in **Supplementary Table 6**): PD-1 BV421, CD20 BV510, CD14 BV510, CD27 BV605, CD8 BV650, CCR6 BV786, CD45RA R718, CD4 APC-H7, CXCR3 Pe-Dazzle594, CXCR5 PeCy7, CD49b FITC, CCR5 RB613, ICOS-PerCP-Cy5.5, CD127 RB744, CD71 RB780, CXCR6 BUV395, CD25 BUV496, CD38 BUV737, CD3 BUV805, and Live/Dead Aqua. PBMC were incubated with 50nM dasatinib for 30 minutes at 37C prior to the addition of pHLA tetramers for an additional 60 minutes at 37C. Cells were then washed in PBS and stained with Live/Dead amine reactive dye for 3 minutes at room temperature, quenched with FACS buffer and incubated with the surface antibody cocktail for 30 minutes at 4C. Following this, cells were washed twice in FACS buffer, and fixed in 1% paraformaldehyde prior to acquisition on a BD Symphony SE flow cytometer.

Analysis of freshly collected PBMC and LN cell suspensions from the FNB cohort was performed with the following antibodies for flow cytometry (details in **Supplementary Table 6**): CD45RA FITC, ICOS PerCP-Cy5.5, CD4 AlexaFluor700, CD20 APC-Fire750, CD3 BUV395, CD19 BUV496, CXCR5 BUV737, PD-1 BV421, CD14 BV510, CXCR3 BV605, CD95 BV650, CD38 BV786, CCR6 Pe-Dazzle594, CD71 PeCy5, IgD PeCy7, and Live/Dead Aqua. Likewise, these samples were analysed using TotalSeqC antibodies (details in **Supplementary Table 7**): CXCR5 (C0144), PD-1 (C0088), CD57 (C0168), ICOS (C0171), TIGIT/VSTM3 (C0089), CXCR3 (C0140), CCR6 (C0143), CCR4 (C0071), CD38 (C0410), CD27 (C0154), CCR7 (C1381), CCR5 (C0141), LAG-3/CD223 (C0152), CD39 (C0176), CD10 (C0062), IgG Fc (C0375), CD95 (C0156), IgM (C0136), CD138 (C0831), CD71 (C0394), and CD69 (C0146). Fresh PBMC and LN samples were incubated with with 50nM dasatinib for 30 minutes at 37C prior to the addition of pHLA tetramers. After 30 minutes, chemokine receptor antibodies (CXCR3, CXCR5, CCR6) were added to the staining cocktail. After an additional 30 minutes incubation, cells were washed in PBS and stained with Live/Dead amine reactive dye for 3 minutes. This was followed by a cocktail of B cell probes and all remaining flow cytometry and TotalSeq antibodies in FACS buffer. Each participant and tissue sample were barcoded with a distinct anti-human TotalSeqC hashtag (BioLegend). Unfixed samples were resuspended in FACS buffer, filtered through a 70um filter, and sorted on a BD FACS Aria III with a 100um filter at 20psi.

#### Activation Induced Marker (AIM) Assay

Analysis of longitudinal PBMC samples by AIM assay was performed using the following antibodies (details in **Supplementary Table 6**): CD154 BUV496, CD137 BV421, CD20 BV510, CD14 BV510, CD4 BV605, CD8 BV650, CCR6 BV786, CXCR4 APC, CD69 APC-Fire750, CXCR5 PE, CXCR3 PE-Dazzle594, CD45RA FITC, OX-40 PerCP-Cy5.5 and Live/Dead Aqua. Longitudinal AIM assays were performed for a subset of participants (n=10) where sufficient PBMC were available. Cryopreserved PBMCs were thawed in RF10, seeded at 1 to 2.5 × 10^6^ cells per well in a 96-well plate, and rested for 3-4 hours at 37°C. The cells were stimulated with peptide pools of 15mers overlapping by 11 amino acids for YFV-17D E, C or NS3 (1 μg/ml, Genscript), or an equivalent volume of DMSO for 20 hours. Each experiment included a positive control well stimulated with SEB (5 μg/ml). CD154 BUV496 antibody was included in the culture medium for the duration of the stimulation. The cells were then washed in PBS and stained with Live/Dead Aqua and surface stained with the remaining antibodies listed above. Cells were washed twice in FACS buffer and fixed in 1% paraformaldehyde prior to acquisition on a BD Symphony SE flow cytometer. AIM^+^ cells were defined as CXCR4^-^CD137^+^ as previously described^40^.

#### Immunophenotyping of B and T cells – Murine samples

Mouse LN suspensions were stained immediately after processing. Half of the total LN cells were resuspended in 50nM dasatinib and incubated for 30 minutes at 37C prior to the addition of IA^b^ tetramers. After a further 60 minutes, cells were washed in FACS buffer and stained as above with Live/Dead Red and the T cell surface antibody cocktail (details in **Supplementary Table 8**): CXCR5 BV421, CD45.2 V500, CD25 BV605, CD4 BV650, PD-1 BV785, TIGIT APC, CXCR6 APC-Fire750, F4/80 Pe-Dazzle594, CD127 Spark PLUS B488, CD69 RB613, CD44 BB700, CCR5 RY775, CD62L BUV395, CD90 BUV496, CD3 BUV737, and B220 BUV805.

The remaining LN cells were stained as above with Live/Dead Red, YFV-17D E B cell probes and the B cell surface antibody cocktail: CD45.2 V500, B220 BUV805, CD3 BUV737, F4/80 Pe-Dazzle594, CD138 BV605, CD38 APC-Fire750, CD98 PeCy7, GL7 AlexaFluor488, and IgD BUV395.

### Generation of sequencing libraries

Sorted T and B cell populations from LN and PBMC samples were processed using 10x Genomics Next GEM 5’ kit v4 (PN-1000265), Next GEM Chip K Single Cell kit (PN-1000286), TCR Amplification Kit (PN-1000252), BCR Amplification kit (PN-1000253) and Dual Index kit TT Set A (PN-1000215). Gene expression, ADT, and T or B cell V(D)J libraries were sequenced on a NovaSeq 6000 (Illumina), targeting a median sequencing depth of 30k, 5k and 7.5k read pairs per cell, respectively.

### Processing and analysis of single-cell gene expression and ADT

Pair-end de-multiplexed FASTQ reads were processed using Cell Ranger v.6.0.1 multi pipeline with reference to GRCh38 human reference genome using default parameters. Quality control was performed in Rstudio (2026.07.1+147) using Seurat v5.5.1. Briefly, UMI counts of gene expression were log-normalized. Quality control was performed by removal of cells with excessive mitochondrial content (5%) or with low or excessive number of features per UMI. ADT counts were normalized using a centered log ratio transformation. Datasets from each FNB experiment were integrated using CCA integration. Highly variable genes were identified using the “FindVariableFeatures” function after removal of immunoglobulin and T cell receptor genes. The data were scaled and centred, and principal component analysis (PCA) performed based on variable gene expression. Cells were clustered using the FindNeighbors and FindClusters functions, and embedded in Uniform Manifold Approximation and Projections (UMAP). Cluster identities were assigned based on differential expression of canonical marker genes for different B or T cell subsets. Additional filtering was undertaken to remove debris or cells expressing markers inconsistent with lymphocytes yielding final datasets of 22150 B cells and 29707 T cells. Gene annotation and clonotypic relationships of BCR and TCR receptor sequences assembled by Cell Ranger were analysed using scRepertoire v2.8.0 (ref).

### Sequencing of single tetramer-positive T cells and E-specific memory B cells

For T cell sorting, cryopreserved PBMC were stained with pHLA tetramers conjugated to SaV-PE, -APC, -BB515 or -BV711 and Live/Dead Aqua, CD20 BV510, CD14 BV510, CD3 APC-H7, CD4 BV605, CD8 BV650 and CD45RA R718 as above. For B cell sorting, samples were stained with PE- and APC-labelled E probes and Live/Dead Aqua, CD3 BV510, CD14 BV510, CD8a BV510, CD16 BV510, CD19 ECD, IgD PeCy7, and IgG BV786. After washing, cells were sorted using a BD Aria III into 96-well plates, subject to cDNA generation and multiplex PCR and Sanger sequencing, as previously described^41,42^.

### Monoclonal antibody generation and expression

Immunoglobulin sequences recovered from 10x transcriptomic pipelines or single cell sequencing were further annotated using ImMunoGeneTics (IMGT) / HighV-Quest (https://www.imgt.org/HighV-QUEST/^43^). The frequency of somatic mutations for a given V-gene sequence were determined by quantifying nonsynonymous nucleotide substitutions with reference to the closest matching germline within the IMGT database. Unmutated common ancestor (UCA) sequences were generated using inferred reversions to nucleotides encoded in the closest matched germline. Clonal inference was performed based on paired heavy and light chains from the same samples. Genes encoding heavy and light chains of antibodies of interest were synthesised (IDT-DNA) and cloned into mammalian expression vectors utilising a human IgG1 Fc backbone. Heavy and light chain plasmids were co-transfected into either Expi293 or ExpiCHO cells (ThermoFisher) and purified from culture supernatants using Protein-A affinity chromatography (Pierce).

## Statistical Analyses

Data are generally presented as median +/- interquartile range. Correlations were analysed using Spearman’s rank-order tests. All statistical analyses were performed using Prism v11 (GraphPad).

## Data availability

10x Genomics raw sequencing data have been deposited at Sequence Read Archive and Gene Expression Omnibus under BioProject XXXX (TBD). Processed transcriptomics and immune receptor data and coding have been deposited at Zenodo at XXXX (TBD). All other data are available from the corresponding authors upon reasonable request.

## Acknowledgements

The authors thank the study participants for providing samples, Kate Merlin and the Clinical Trials and Biorepository Laboratory Team at St Vincent’s Centre for Applied Medical Research for phlebotomy and sample processing, and Eric Lam for assistance with cell sorting. Transcriptomics and Sequencing Services for this study were provided by the Garvan Genomics Platform at the Garvan Institute of Medical Research. This study was funded by NHMRC grants to JAJ, AKW and CMLM (GNT2009308, GNT2026762, GNT2012863). WSL, SJK, ADK, JAJ and AKW are supported by NHMRC Investigator grants. JAJ is supported by the Sylvia and Charles Viertel Charitable Foundation. RBE is supported by the Elizabeth and Vernon Puzey Scholarship and the Research Training Program Scholarship.

## Authors’ contributions

RBE, WSL, TGP, SJK, ADK, CMLM, JAJ and AKW designed the study. RBE, WSL, PM, CAG, MZMZ, RE, ER, KMW, AK, JZ, SO, BM, CMLM, JAJ and AKW performed experiments. SB, NM and DW contributed unique reagents. RBE, WSL, PM, CAG, JJZ, SJK, ADK, CMLM, JAJ and AKW analysed the experimental data. RBE, WSL, CAG, CMLM, JAJ and AKW wrote the manuscript. All authors reviewed the manuscript.

---

**Supplementary Table 1.** Cohort demographics.

| Participant | Age | Gender | Tissues Sampled | Relevant HLA-DR alleles |
|---|---|---|---|---|
| CP01 | 22 | F | Blood | 03:01, 11:01 |
| CP46 | 53 | M | Blood | 03:01, 07:01 |
| CP47 | 57 | F | Blood | 3:01 |
| CP53 | 29 | F | Blood | 03:01, 07:01 |
| CP55 | 25 | M | Blood | 03:01, 07:01 |
| CP57 | 58 | M | Blood | 3:01 |
| CP60 | 50 | F | Blood | 03:01, 15:01 |
| COR006 | 28 | F | Blood | 04:01, 11:01 |
| COR004 | 29 | M | Blood | N/A |
| COR299 | 43 | F | Blood | 11:01 |
| COR414 | 42 | M | Blood | N/A |
| COR396 | 35 | M | Blood, dLN | 04:01, 11:01 |
| COR418 | 24 | F | Blood, dLN | 3:01 |
| COR282 | 34 | M | Blood, dLN | 1:01 |
| COR424 | 43 | M | Blood, dLN | 03:01, 04:01 |
| COR429 | 32 | F | Blood, dLN, ndLN | N/A |
| COR430 | 27 | F | Blood, dLN, ndLN | 15:01 |

**Supplementary Table 6.** Antibodies for human clinical samples.

| Antigen | Fluorochrome | Clone | Source | Dilution | RRID |
|---|---|---|---|---|---|
| CD11c | RB613 | B-ly6 | Waters | 1:50 | AB_3686242 |
| IgA | VioBlue | IS11-8E10 | Miltenyi | 1:33 | AB_2733959 |
| CD14 | BV510 | M5E2 | Waters | 1:100 | AB_2739916 |
| CD3 | BV510 | OKT3 | BioLegend | 1:400 | AB_2563704 |
| CD8a | BV510 | RPA-T8 | BioLegend | 1:1000 | AB_2561942 |
| CD16 | BV510 | 3G8 | BioLegend | 1:300 | AB_2562085 |
| CD27 | BV605 | O323 | BioLegend | 1:25 | AB_2561450 |
| IgG | BV786 | G18-145 | Waters | 1:50 | AB_2738684 |
| CD38 | AlexaFluor700 | HIT2 | BioLegend | 1:100 | AB_2072781 |
| CD20 | APC-Fire750 | 2H7 | BioLegend | 1:100 | AB_2572126 _ |
| CD19 | ECD | J3-119 | Beckman Coulter | 1:100 | AB_130854 |
| CD71 | PeCy5 | CY1G4 | BioLegend | 1:200 | AB_2936703 |
| IgD | PeCy7 | IA6-2 | Waters | 1:200 | AB_10642457 |
| IgM | BUV395 | G20-127 | Waters | 1:100 | AB_2721269 |
| CD21 | BUV737 | B-ly4 | Waters | 1:100 | AB_2870115 |
| PD-1 | BV421 | EH12.2H7 | Waters | 1:50 | AB_11153482 |
| CD20 | BV510 | 2H7 | Waters | 1:100 | AB_2737985 |
| CD8 | BV650 | RPA-T8 | BioLegend | 1:400 | AB_2563505 |
| CCR6 | BV786 | G034E3 | BioLegend | 1:100 | AB_2563660 |
| CD45RA | R718 | HI100 | Waters | 1:100 | AB_2916420 |
| CD4 | APC-H7 | RPA-T4 | Waters | 1:50 | AB_1645478 |
| CXCR3 | Pe-Dazzle594 | G02H57 | BioLegend | 1:50 | AB_2564288 |
| CXCR5 | PeCy7 | MU5UBEE | ThermoFisher | 1:33 | AB_2573540 |
| CD49b | FITC | AK-7 | Waters | 1:20 | AB_395888 |
| CCR5 | RB613 | 2D7/CCR5 | Waters | 1:50 | AB_3691403 |
| ICOS | PerCP-Cy5.5 | C398.4A | BioLegend | 1:50 | AB_10641280 |
| CD127 | RB744 | HIL-7R-M21 | Waters | 1:100 | AB_3685886 |
| CD71 | RB780 | M-A712 | Waters | 1:400 | AB_3688144 |
| CXCR6 | BUV395 | 13B | Waters | 1:50 | AB_2741615 |
| CD25 | BUV496 | 2A3 | Waters | 1:100 | AB_2870761 |
| CD38 | BUV737 | HB7 | Waters | 1:67 | AB_2870148 |
| CD3 | BUV805 | SK7 | Waters | 1:100 | AB_2870181 |
| CD45RA | FITC | HI100 | BioLegend | 1:50 | AB_314410 |
| CD4 | AlexaFluor700 | RPA-T4 | Waters | 1:100 | AB_396943 |
| CD3 | BUV395 | SK7 | Waters | 1:200 | AB_2744382 |
| CD19 | BUV496 | SJ25C1 | Waters | 1:400 | AB_2870221 |
| CXCR5 | BUV737 | MU5UBEE | ThermoFisher | 1:50 | AB_3074035 |
| CXCR3 | BV605 | G02H57 | BioLegend | 1:50 | AB_2563157 |
| CD95 | BV650 | DX2 | BioLegend | 1:50 | AB_2632622 |
| CD38 | BV786 | HIT2 | BioLegend | 1:50 | AB_2565893 |
| CCR6 | PeDazzle-594 | G034E3 | BioLegend | 1:50 | AB_2564233 |
| CD154 | BUV496 | TRAP1 | Waters | 1:100 | AB_2874577 |
| CD137 | BV421 | 4B4-1 | BioLegend | 1:50 | AB_2563830 |
| CD4 | BV605 | RPA-T4 | BioLegend | 1:100 | AB_2564391 |
| CXCR4 | APC | 12G5 | Waters | 1:50 | AB_398616 |
| CD69 | APC-Fire750 | FN50 | BioLegend | 1:50 | AB_2616709 |
| CXCR5 | PE | MU5UBEE | ThermoFisher | 1:50 | AB_11219877 |
| OX-40 | PerCP-Cy5.5 | ACT35 | BioLegend | 1:50 | AB_10719224 |
| CD3 | APC-H7 | SK7 | Waters | 1:100 | AB_1645475 |

**Supplementary Table 7.** Antibodies used for scRNAseq.

| **Antigen** | **Company** |  | **RRID** |
|---|---|---|---|
| CXCR5 | BioLegend | C0144 | AB_2800968 |
| PD-1 | BioLegend | C0088 | AB_2800862 |
| CD57 | BioLegend | C0168 | AB_2801030 |
| ICOS | BioLegend | C0171 | AB_2800823 |
| TIGIT/VSTM3 | BioLegend | C0089 | AB_2801021 |
| CXCR3 | BioLegend | C0140 | AB_2800949 |
| CCR6 | BioLegend | C0143 | AB_2810563 |
| CCR4 | BioLegend | C0071 | AB_2800988 |
| CD38 | BioLegend | C0410 | AB_2820007 |
| CD27 | BioLegend | C0154 | AB_2800747 |
| CCR7 | BioLegend | C1381 | AB_2936612 |
| CCR5 | BioLegend | C0141 | AB_2810570 |
| LAG-3/CD223 | BioLegend | C0152 | AB_2814327 |
| CD39 | BioLegend | C0176 | AB_2800853 |
| CD10 | BioLegend | C0062 | AB_2800817 |
| IgG Fc | BioLegend | C0375 | AB_2801087 |
| CD95 | BioLegend | C0156 | AB_2800787 |
| IgM | BioLegend | C0136 | AB_2800835 |
| CD138 | BioLegend | C0831 | AB_2814282 |
| CD71 | BioLegend | C0394 | AB_2800885 |
| CD69 | BioLegend | C0146 | AB_2800810 |
| Hashtag 1 (CD298/b2M) | BioLegend | C0251 | AB_2801031 |
| Hashtag 2 (CD298/b2M) | BioLegend | C0252 | AB_2801032 |
| Hashtag 3 (CD298/b2M) | BioLegend | C0253 | AB_2801033 |
| Hashtag 4 (CD298/b2M) | BioLegend | C0254 | AB_2801034 |

**Supplementary Table 8.** Antibodies for mouse tissue samples.

| **Antigen** | **Fluorochrome** | **Clone** | **Company** | **Dilution** | **RRID** |
|---|---|---|---|---|---|
| CXCR5 | BV421 | L138D7 | BioLegend | 1:50 | AB_2562128 |
| CD45.2 | V500 | 104 | Waters | 1:200 | AB_10897142 |
| CD25 | BV605 | PC61 | Waters | 1:100 | AB_2737982 |
| CD4 | BV650 | RM4-5 | Waters | 1:300 | AB_2716859 |
| PD-1 | BV785 | 29F.1A12 | Waters | 1:100 | AB_3684376 |
| TIGIT | APC | 4D4/mTIGIT | BioLegend | 1:33 | AB_2750515 |
| CXCR6 | APC-Fire750 | SA051D1 | BioLegend | 1:50 | AB_2910304 |
| F4/80 | Pe-Dazzle594 | BM8 | BioLegend | 1:200 | AB_2564133 |
| CD127 | Spark PLUS B488 | A7R34 | BioLegend | 1:67 | AB_3720645 |
| CD69 | RB613 | H1.2F3 | Waters | 1:50 | AB_3690381 |
| CD44 | BB700 | IM7 | Waters | 1:1000 | AB_2744396 |
| CCR5 | RY775 | C34-3448 | Waters | 1:33 | AB_3692682 |
| CD62L | BUV395 | MEL-14 | Waters | 1:400 | AB_3685037 |
| CD90 | BUV496 | 53-2.1 | Waters | 1:200 | AB_2870661 |
| CD3 | BUV737 | 145-2C11 | Waters | 1:100 | AB_2870100 |
| B220 | BUV805 | RA3-6B2 | Waters | 1:140 | AB_3684861 |
| CD138 | BV605 | 281.2 | Waters | 1:200 | AB_2721029 |
| CD38 | APC-Fire750 | 90 | BioLegend | 1:200 | AB_2876402 |
| CD98 | PeCy7 | RL388 | BioLegend | 1:1000 | AB_2750546 |
| GL7 | AlexaFluor488 | GL7 | BioLegend | 1:3000 | AB_2563285 |
| IgD | BUV395 | 11-26c.2a | Waters | 1:140 | AB_2738723 |

**Extended Data Fig 1.**
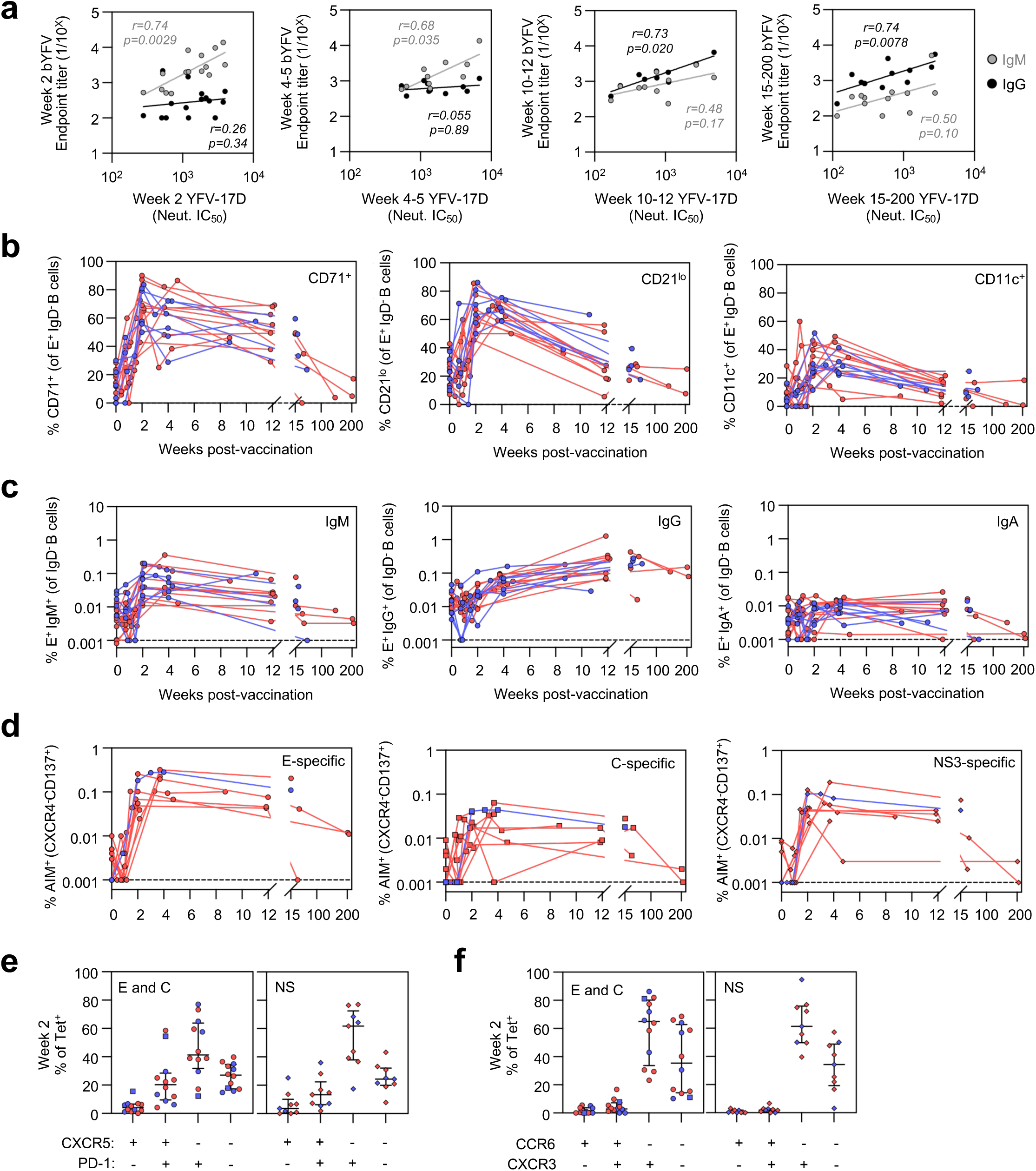
Immune Dynamics of YFV-17D immunization in the blood. **(a)** Association between bYFV virus-like particle IgM (grey) or IgG (black) ELISA endpoint titers and neutralizing antibody titres at weeks 2, 4-5, 10-12 or 15-200. N=17, Spearman r and p values indicated. Longitudinal frequency of E-specific **(b)** CD71^+^, CD21^lo^ and CD11c^+^ or **(c)** IgM^+^, IgG^+^ and IgA^+^ B cells (N=17). **(d)** Frequency of AIM^+^ (CXCR4^-^CD137^+^) CD45RA^-^CD4^+^CD3^+^ T cells following stimulation with E, C or NS3 peptide pools (N=10). **(e)** Expression of CXCR5 and PD-1 or **(f)** CCR6 and CXCR3 by Tet^+^ cells at week 2 post-vaccination, with specificity for E or C (left panel) and NS epitopes (right panel). Lines indicate median and IQR (N=13). Red, participants with blood sampling only; blue, participants with blood and FNB sampling.

**Extended Data Fig 2.**
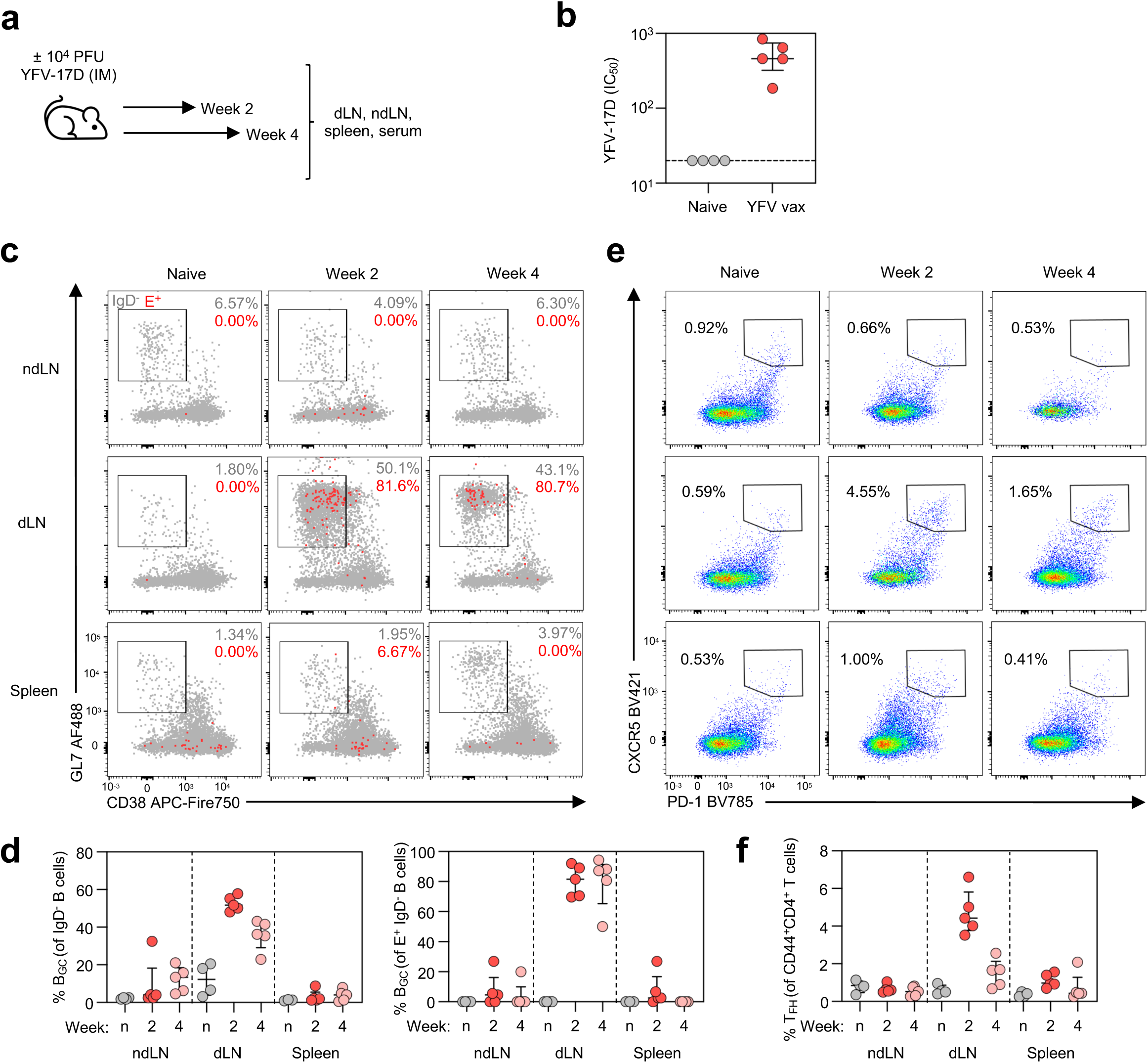
YFV-17D immunity is programmed in the vaccine draining lymph node in a mouse model. **(a)** Mice were vaccinated intramuscularly (IM) with 10^4^ PFU of live YFV-17D virus or mock vaccinated (naïve, n). 2 or 4 weeks later, blood, draining lymph nodes (dLN), non-draining lymph nodes (ndLN) and spleen were collected. **(b)** Antibody titres from a live virus neutralization assay for week 2-post YFV-17D (N=5) or naïve mouse serum (N=4). **(c)** Representative plots and **(d)** frequencies of B_GC_ (GL7^+^CD38^-^) among total IgD^-^ B cells (CD45^+^CD3^-^B220^+^IgD^-^) or E^+^ IgD^-^ B cells. **(e)** Representative plots and **(f)** frequencies of T_FH_ (PD-1^++^CXCR5^++^) among CD44^+^ CD4^+^ T cells (B220^-^ CD45^+^CD3^+^CD4^+^CD44^+^). N=18 mice (4-5 per group). Lines indicate median and IQR.

**Extended Data Figure 3.**
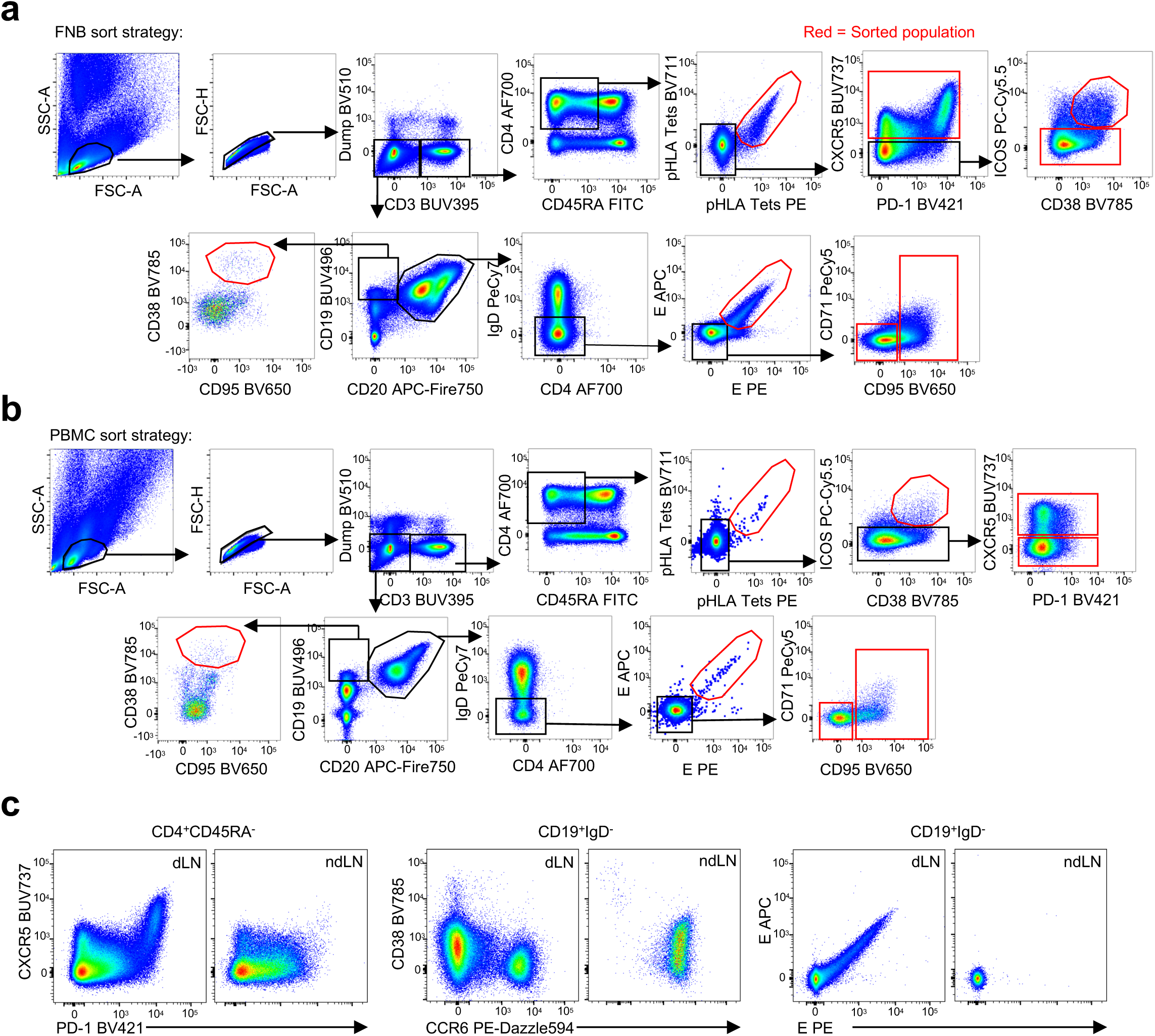
B cell and CD4^+^ T cell populations sorted for transcriptional analysis. Sort strategy for freshly isolated **(a)** FNB and **(b)** PBMC samples. Representative plots show matched PBMC and LN samples from a single participant at week 4 post-vaccination. Red gates indicate populations sorted for downstream capture. **(c)** Enrichment of active GC responses in draining (dLN, ipsilateral) vs non-draining (ndLN, contralateral) axillary LN at week 4.

**Extended Data Figure 4.**
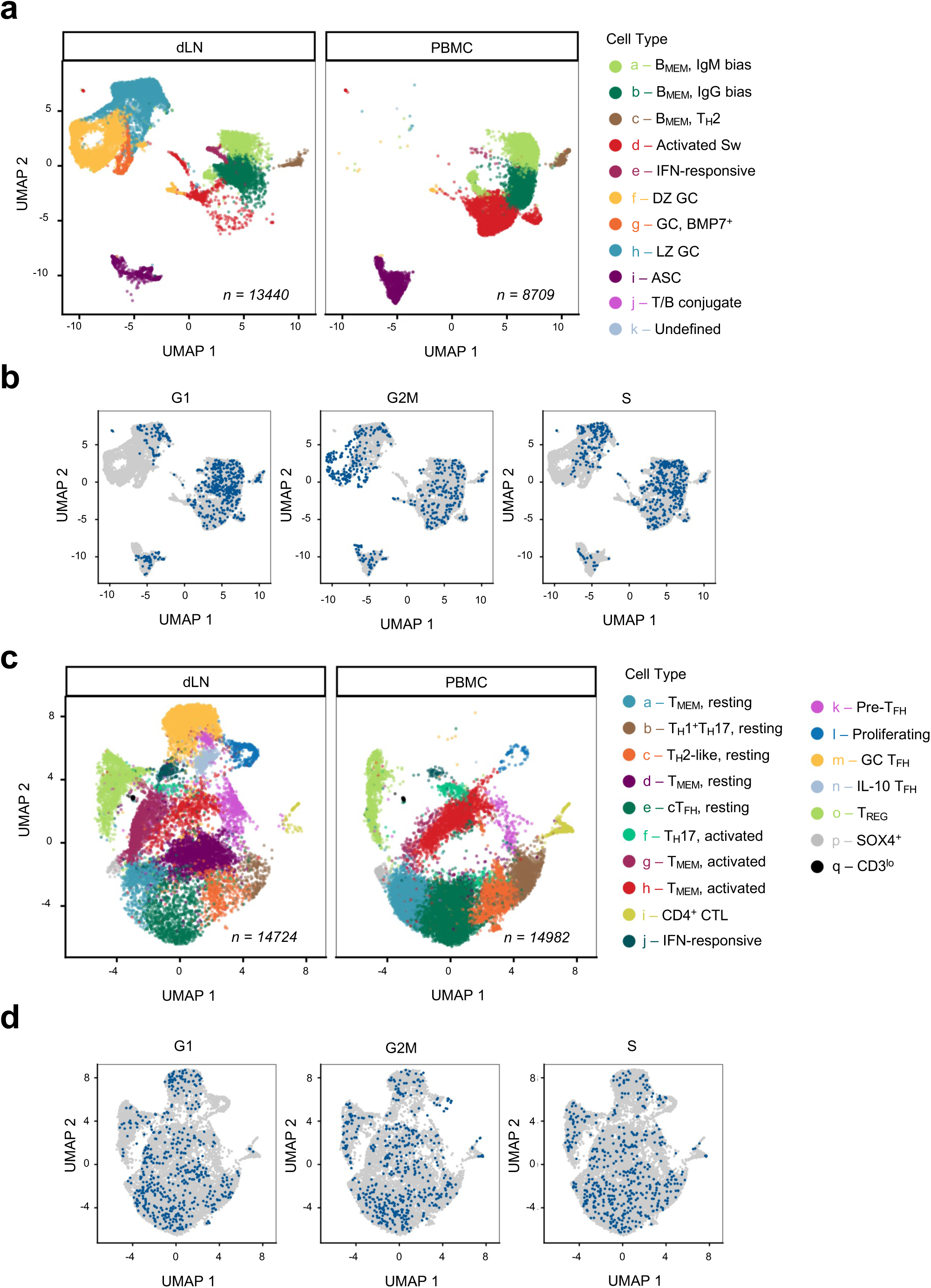
Tissue distribution and cell cycle of sorted B cell and CD4^+^ T cell clusters. (a, **c)** UMAP displaying all sorted B cells or CD4^+^ T cells at weeks 2 and 4 (N=6 participants), segregated by tissue-specific hashtags. **(b, d)** Cell cycle analysis highlighting cells in G1, G2M and S phase (blue) across all tissues and timepoints.

**Extended Data Figure 5.**
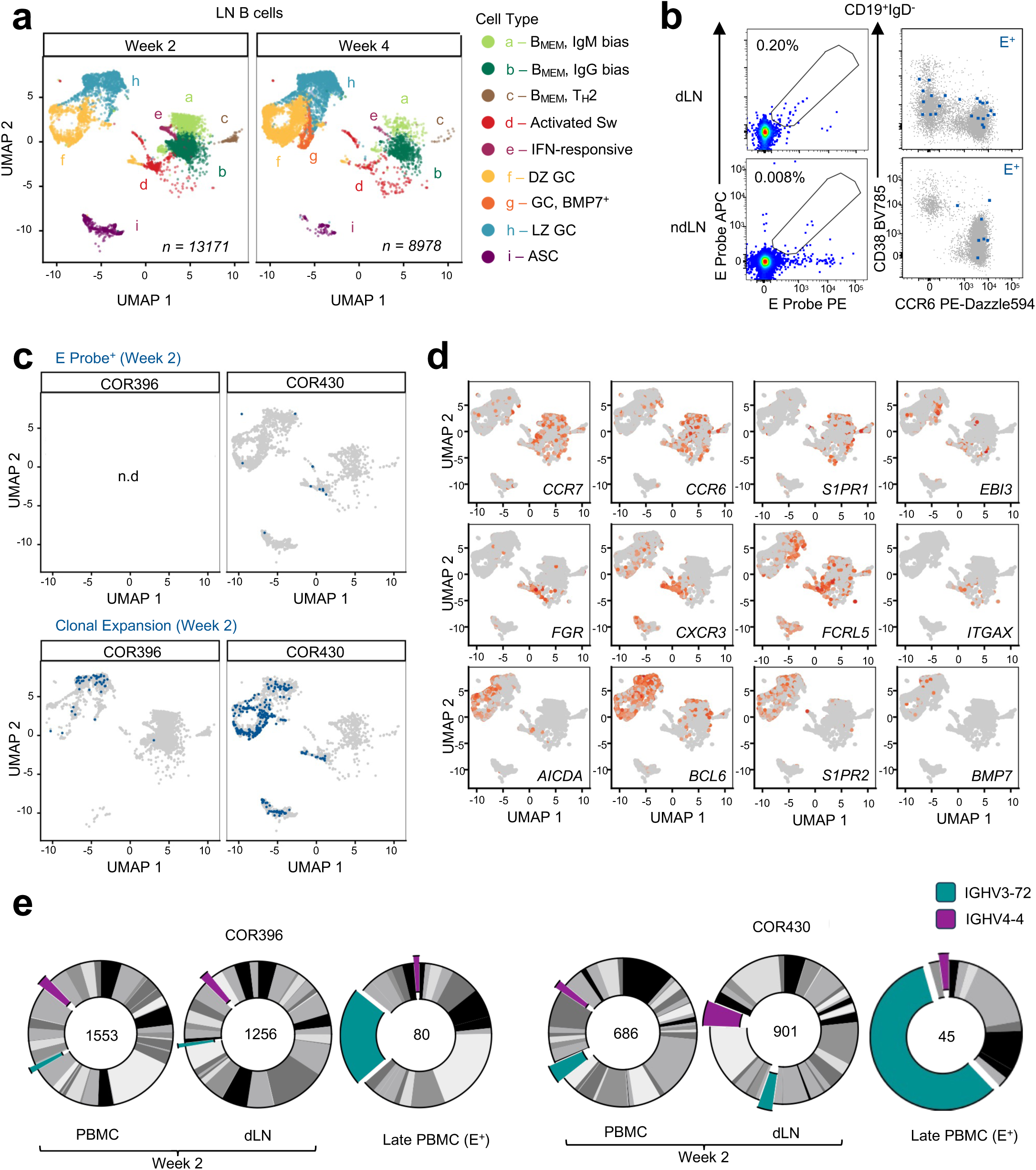
Lymph node B cell populations at week 2 post-vaccination. **(a)** UMAP clustering of draining lymph node B cells captured by FNB at weeks 2 (N=2) and 4 (N=3). **(b)** Staining of E probe^+^ class-switched B cells at week 2 in draining and non-draining (contralateral) LN from COR430. Blue, E-specific cells; grey, CD19^+^IgD^-^ cells. **(c)** Overlay of E probe^+^ or clonally expanded (5+ clonotypes) B cells (blue) at week 2 for participants COR396 and COR430. **(d)** Log-normalised expression of selected genes across clusters at week 2. **(e)** Distribution of IGHV gene utilisation for immunoglobulins recovered from sorted PBMC or LN B cells at week 2 post immunisation, or from E-specific memory B cells recovered in the blood at weeks 9 (COR430) or 53 (COR396). Proportions of IGHV3-72 and IGHV4-4 utilisation are indicated in colour.

**Extended Data Figure 6.**
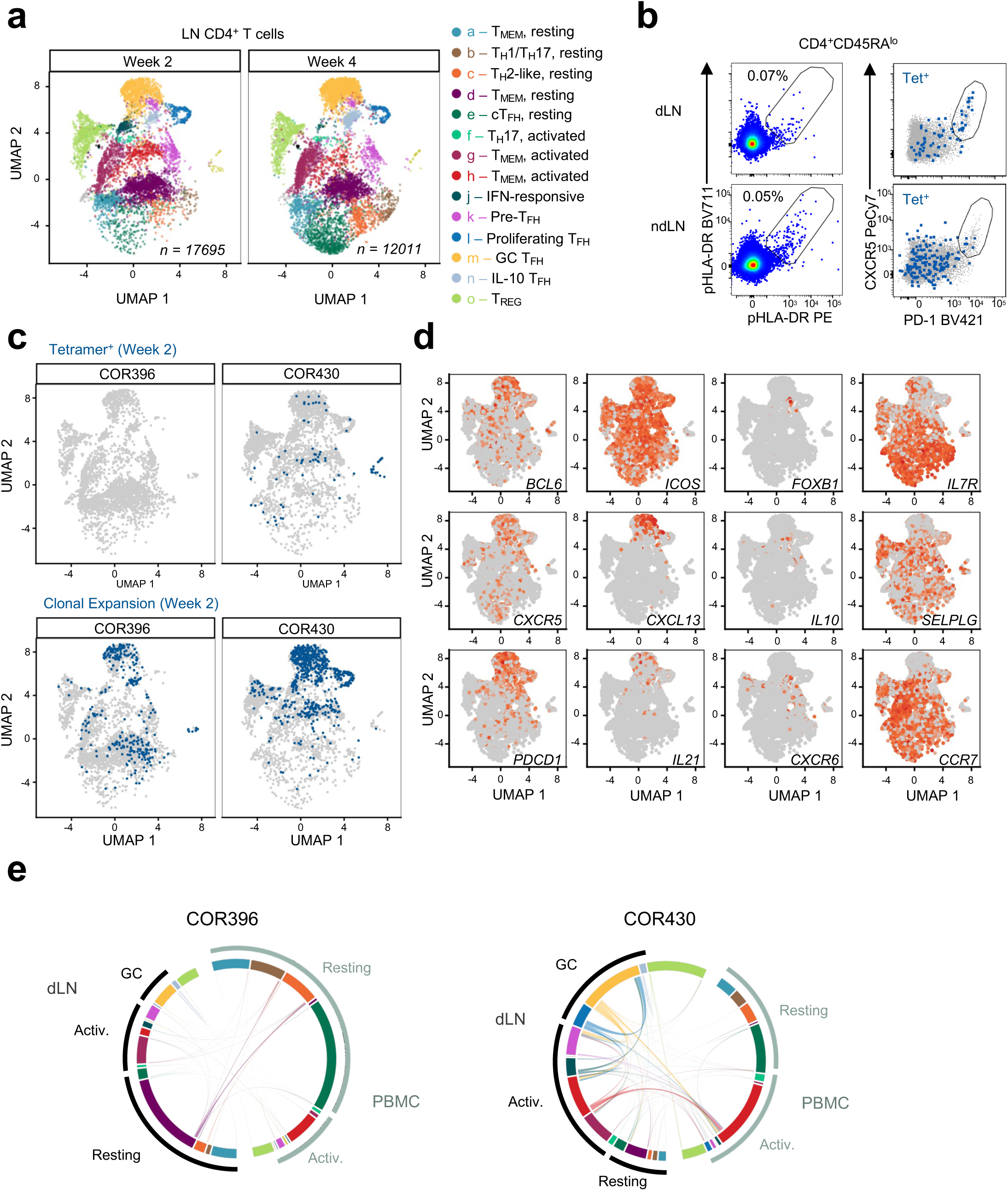
Lymph node T cell populations at week 2 post-vaccination.(a) UMAP clustering of draining lymph node CD4^+^ T cells captured by FNB at week 2 (N=2) and week 4 (N=3). **(b)** Staining of pHLA tetramers in draining and non-draining (contralateral) LN from COR430 (week 2). Blue, tetramer^+^ cells; grey, bulk CD4^+^CD45RA^-^ cells. **(c)** UMAP overlay of tetramer+ or clonally expanded CD4^+^ T cells (blue) in the dLN for participants COR396 and COR430. **(d)** Log-normalised expression of selected genes within the week 2 dLN dataset. **(e)** Circos plots showing shared clones within and between tissues for COR396 and COR430.

**Extended Data Figure 7.**
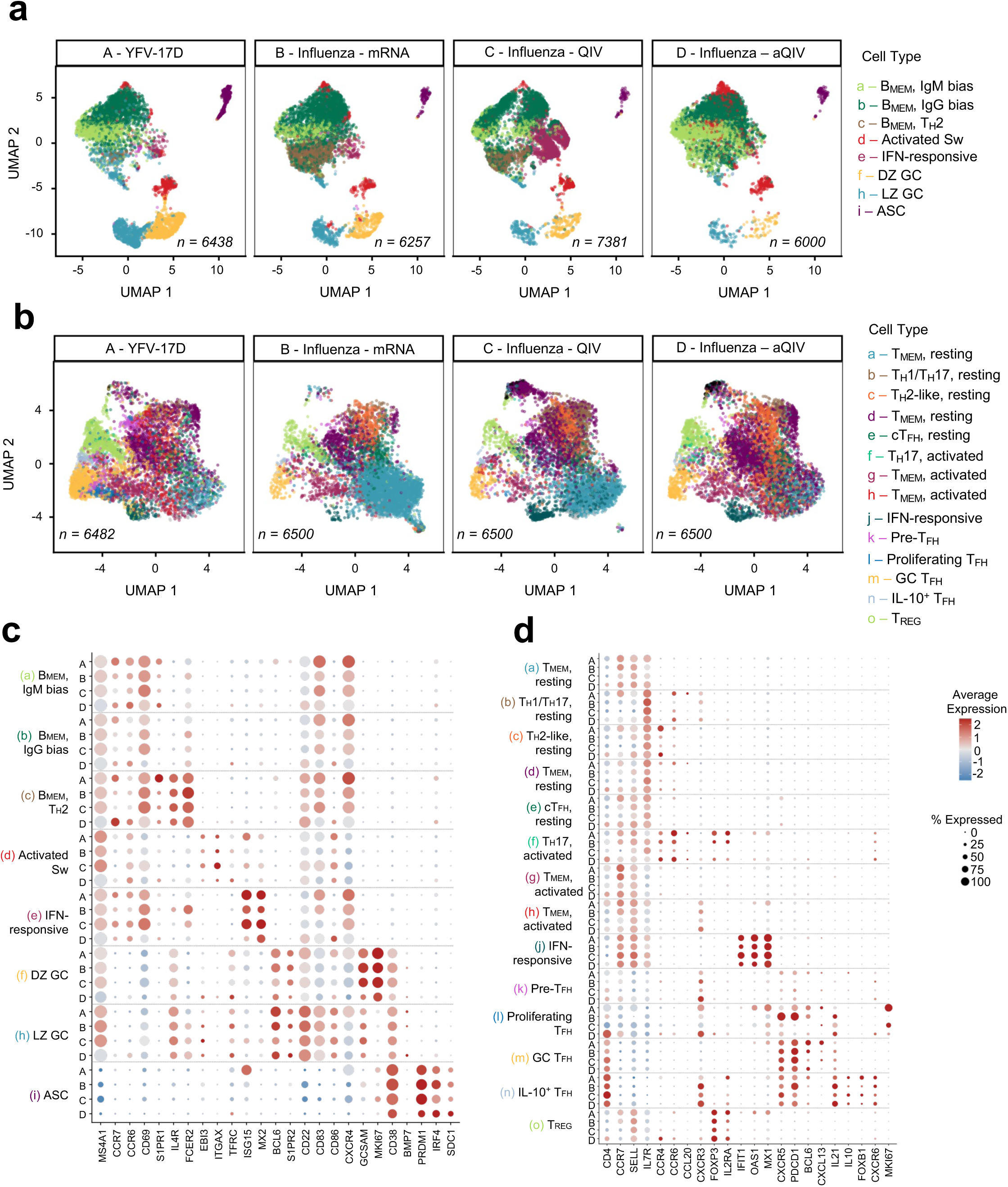
Transcriptional profile of lymph node B and T lymphocyte populations at 2 weeks post-vaccination with YFV-17D versus vaccines for influenza. UMAP representations of lymph node **(a)** B cell and **(b)** CD4 T cell populations observed 2 weeks after immunization with YFV-17D (A; this study), influenza mRNA/LNP (B; Matz et al.^17^), seasonal influenza (C; McIntire et al ^12^ for B cells, Schattgen et al.^10^ for T cells), or adjuvanted seasonal influenza (D; Siu et al^14^) vaccine. Following dataset integration of the different studies, YFV-17D cluster identities from Figure 2A or 3A were projected onto a shared UMAP. Expression of key genes among **(c)** B cell or **(d)** CD4^+^ T cell clusters for each dataset.

**Extended Data Figure 8.**
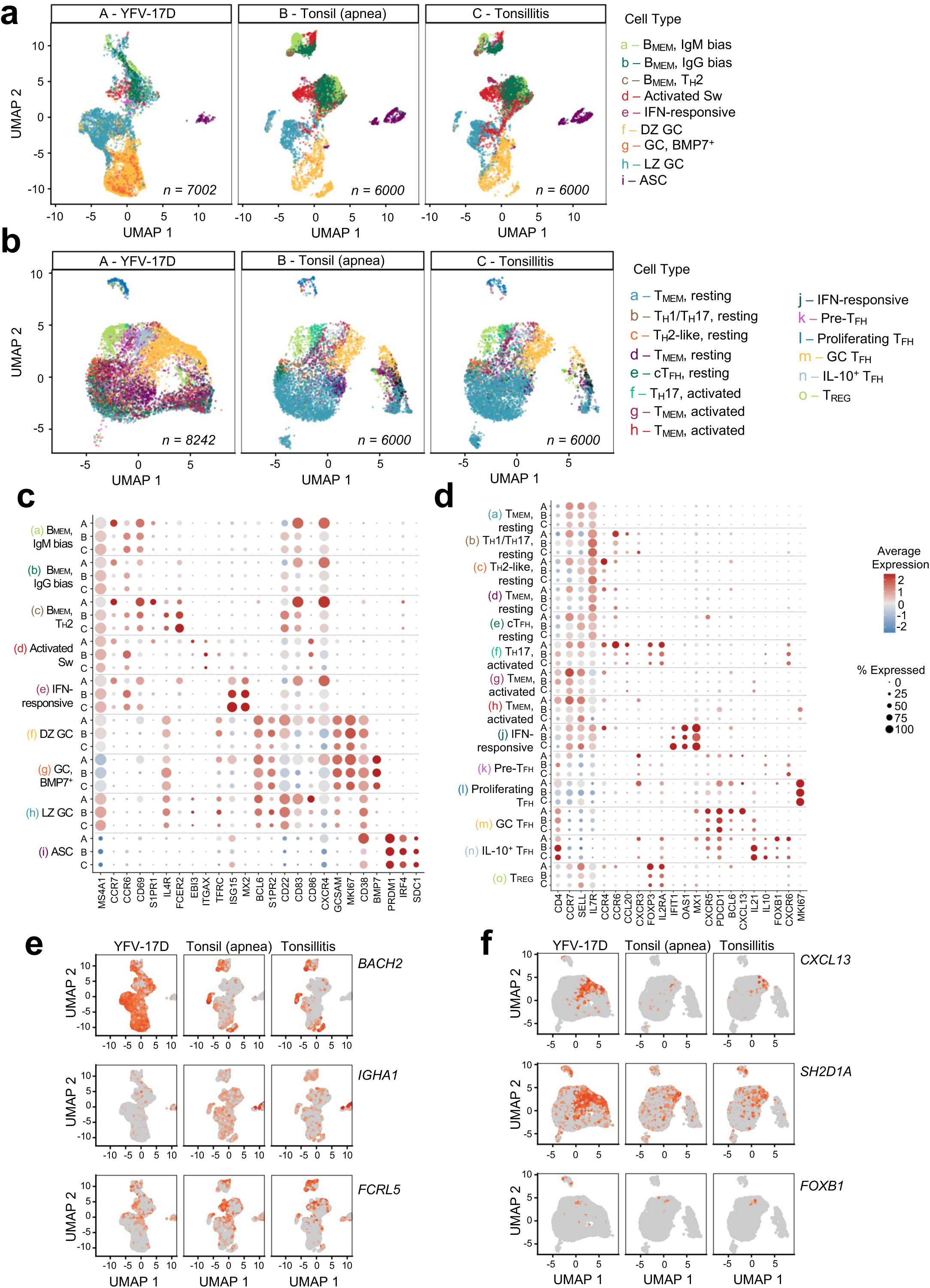
Transcriptional profiles of B and T lymphocyte populations in inflamed tonsils versus post-YFV-17D lymph nodes. UMAP representations of **(a)** B cell and **(b)** CD4 T cell populations in the lymph node at 2 or 4 weeks after immunization with YFV-17D (A; this study), or at steady-state in tonsils removed due to sleep apnea or tonsillitis (B and C, respectively; Massoni-Badosa et al.^18^). Following dataset integration of the different studies, YFV-17D cluster identities from Figure 2A or 3A were projected onto a shared UMAP. Expression of key genes within **(c)** B cell or **(d)** CD4^+^ T cell clusters for each dataset. UMAP projections highlighting differential expression of selected **(e)** B cell or **(f)** T_FH_ genes across tissues.

**Supplementary Figure 1.**
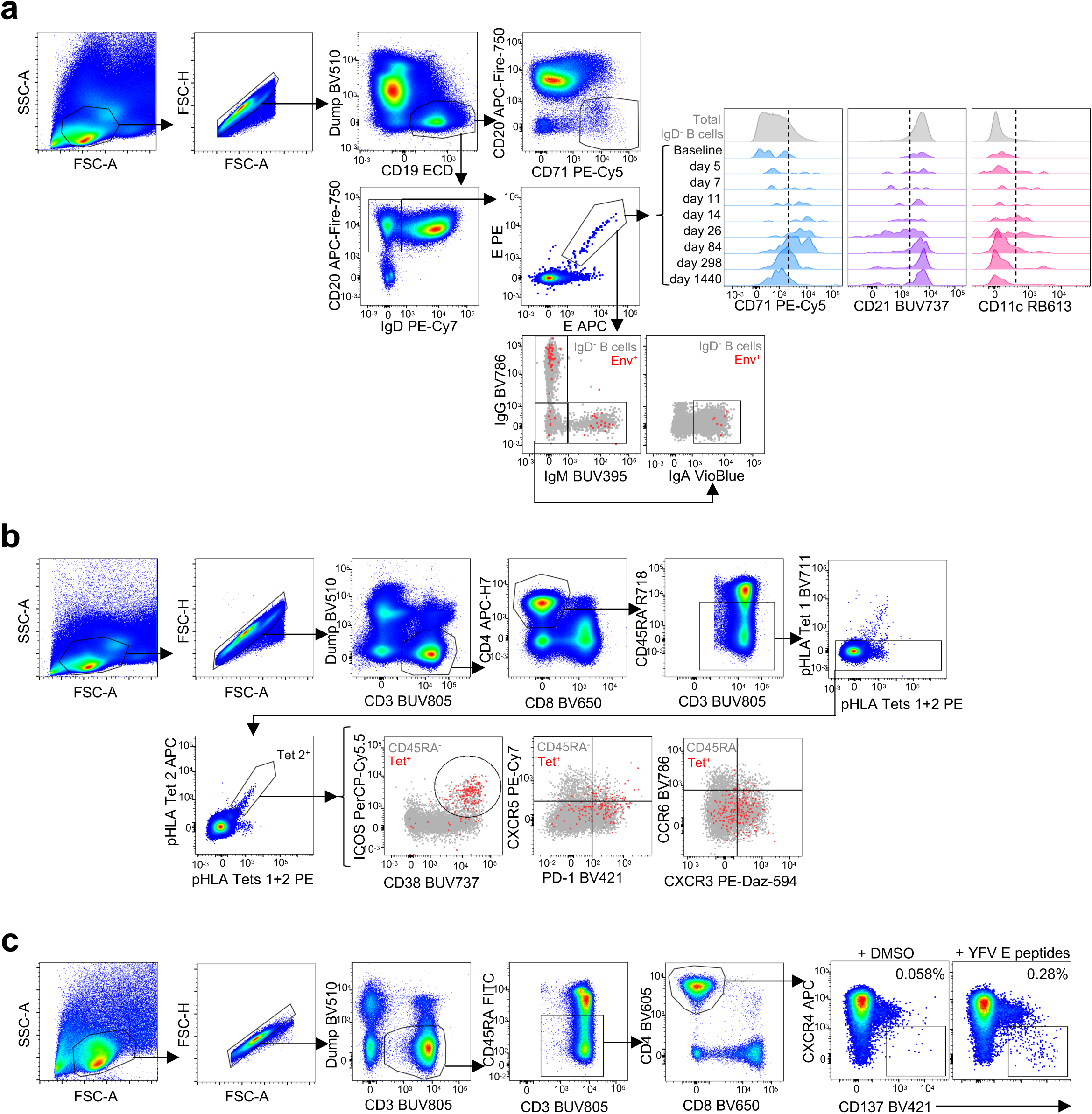
Gating for FACS data shown in Fig 1 and ED Fig 1. **(a)** Representative gating for longitudinal B cell and probe analysis. **(b)** Representative gating for longitudinal CD4^+^ T cell and tetramer analysis. **(c)** Representative gating for longitudinal CD4^+^ T cell AIM analysis.

**Supplementary Figure 2.**
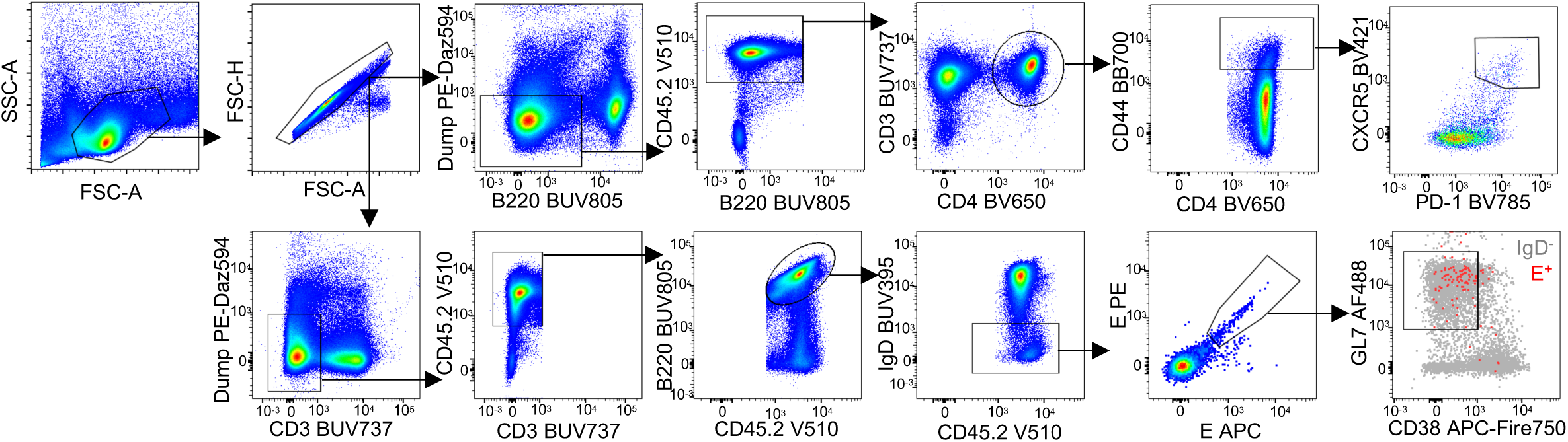
Gating for FACS data shown in ED Figure 2. Gating for longitudinal B cell and CD4^+^ T cell analysis in vaccinated mice.

